# Structural insights into the recruitment of viral Type 2 IRES to ribosomal preinitiation complex for protein synthesis

**DOI:** 10.1101/2025.05.28.656627

**Authors:** Deepakash Das, Tanweer Hussain

**Affiliations:** Department of Developmental Biology and Genetics, Indian Institute of Science, Bengaluru, INDIA. PIN-560012

## Abstract

Picornaviruses employ internal ribosome entry sites (IRESs) in their genomic RNA to hijack the host’s translational machinery. The picornavirus, Encephalomyocarditis virus, employs a type 2 IRES present in its 5’UTR and requires 43S ribosomal preinitiation complex (PIC), the central domain of eukaryotic initiation factor (eIF) 4G, eIF4A, and an essential ITAF (IRES trans-acting factor)-polypyrimidine tract binding protein 1 (PTB1) to form 48S PIC. In this study, we have used cryo-electron microscopy (cryo-EM) to determine the structure of EMCV IRES-bound mammalian 48S PIC in a scanning-arrested closed state at the start codon. The EMCV IRES domains contacts initiator tRNA (tRNA_i_) and 40S head at the inter-subunit interface, which reveals an altogether unique mechanism used by viruses to capture host translational machinery for its protein synthesis. The tRNA_i_ is held away from the 40S body in contrast to canonical cap-dependent translation while the domain I apical region of EMCV IRES mimics 28S rRNA of 60S to interact with 40S ribosomal head proteins-uS13 and uS19. The structural analysis account for numerous biochemical studies on Type 2 IRES and shows how Type 2 IRES interacts with 43S PIC to form 48S PIC. This study provides mechanistic insights for understanding EMCV IRES-mediated translation initiation, which could be extrapolated to other IRESs sharing similar motifs and factor requirements including type 1 viral IRESs.

## Introduction

Eukaryotic translation initiation occurs in two different modes-Cap-dependent and Cap-independent. Cap-dependent translation initiation in eukaryotes can be divided into four major steps-(i) formation of 43S ribosomal PIC [40S subunit in complex with eIF1, eIF1A, eIF3, eIF5 and ternary complex (eIF2-GTP-tRNA_i_)]; (ii) recruitment of 43S PIC to 5’end of capped mRNA, mediated by eIF4F complex (formed by eIF4E,eIF4G and eIF4A) yielding 48S complex (48S PIC); (iii) scanning of 5’ untranslated region (UTR) of mRNA and start codon recognition; and (iv) joining of large 60S ribosomal subunit to form elongation-competent 80S ribosomal complex. Among these, major rate-limiting steps include-the regulation of available tRNA_i_ as the ternary complex (eIF2.GTP.Met-tRNA_i_) and recruitment of mRNA on 43S PIC (*Jackson et al 2010; Brito Querido et al 2024*). Cryo-EM studies on eukaryotic canonical 48S PICs from yeast (*Hussain et al 2014; Llacer et al 2015*) to humans (*Eliseev et al 2018; Simonetti et al 2020; Brito Querido et al 2020; Yi et al 2022; Brito Querido et al 2024; Petrychenko et al 2024*) have revealed various interactions among the initiation factors, mRNA, and ribosome in scanning (P_OUT_ or open) state and scanning-arrested (P_IN_ or closed) states. In mammals, 48S PIC formation is mediated by the interaction of eIF3 and eIF4G (*Villa et al 2013*) and recent attempts to understand the mammalian canonical 48S PIC could capture interactions of eIF4 proteins (eIF4G and eIF4A) with eIF3 (*Brito Querido et al 2020; Brito Querido et al 2024*).

Alternatively, several positive strand RNA viruses use Internal Ribosome Entry Sites (IRESs), which are internal cis-acting sequences present in 5’ UTR of mRNA that drive the assembly of translation initiation complex without the requirement of 7-methylguanosine cap (*Martínez-Salas et al 2018*), initially reported in picornaviruses such as poliovirus (PV) and encephalomyocarditis virus (EMCV) RNA genome (*Jang et al 1988; Pelletier and Sonenberg, 1988; Trono et al 1988*). As of 2020, the IRESbase database reported 1328 IRESs, out of which 554 were viral IRESs from 198 viruses (*Zhao et al 2020*). These RNA elements have unique secondary and tertiary structures, which allows them to hijack the host translational machinery and promote translation initiation internally by recruitment of host ribosome, canonical eIFs, and IRES trans acting factors (ITAFs) via multiple RNA–RNA and RNA–protein interactions (*Lee et al 2017; Martinez-Salas et al 2018*). The IRESs lack sequence homology and enfolds different structural organization, thus requiring different eIFs for the assembly of 48S PICs (*Jackson et al 2010*). The type 1 IRES (represented by PV) and type 2 IRES (EMCV) requires almost all eIFs, except eIF4E or N terminal of eIF4G for the formation of 48S PIC (*Lozano and Martinez-Salas, 2015)*. However, initiation on the latter does not require scanning for start codon recognition (*Kaminski et al 1994; Sweeney et al 2014)*. The type 3 IRES (Hepatitis C virus-HCV) does not require eIF4 factors or scanning for 48S PIC formation (*Niepmann and Gerresheim 2020*). The type 4 IRES (Cricket paralysis virus-CrPV intergenic IRES) initiates without the requirement of any eIFs and tRNA_i_ (*Johnson et al 2017*). The HCV IRES and CrPV intergenic IRES directly interacts with host 40S ribosome, however they differ in their mechanism of attachment. Where, the CrPV IRES (type 4) mimics anticodon-codon interaction via pseudoknot 1 to bind with ribosome in an elongation-competent state (*Petrov et al 2016; Fernandez et al 2014*) the HCV IRES (type 3) binds to the solvent side of 40S subunit by replacing eIF3 and directly interacts with expansion segment (ES7) of 18S rRNA and ribosomal proteins near mRNA exit channel, thus placing the initiation codon at the P-site (*Hashem et al 2013; Niepmann and Gerresheim 2020; Brown et al 2022*). While the structures of ribosome-bound type 4 (*Spahn et al 2004; Fernandez et al 2014; Muhs et al 2015; Murray et al 2016; Pisareva et al 2018; Acosta-Reyes et al 2019; Abaena et al 2020*) and type 3 IRES (*Spahn et al 2004; Boehringer et al 2005; Hashem et al 2013; Yamamoto et al 2014; Quade et al 2015; Yamamoto et al 2015; Yokoyama et al 2019; Brown et al 2022*) have been determined, there is no structural information about Type 2 and Type 1 IRES and their mode of recruitment to 48S PIC.

The 5’UTR of EMCV, genera Cardiovirus and Picornaviridae family (*International Committee on Taxonomy of Viruses Executive Committee, 2020*) folds into various stem-loops numbered D-L, where domains H-L, followed by the initiation codon at the 834^th^ residue (∼450 nucleotides in length) make a functional IRES moiety (*Carocci et al 2012*). It requires the core of 43S PIC, the central domain of eIF4G, eIF4A, and an essential ITAF-polypyrimidine tract binding protein 1 (PTB1), and the presence of eIF4B enhances 48S formation (*Hellen and Wimmer 1995; Martinez-Salaz et al 2018; Sweeney et al 2014*). Functional characterization of these domains showed H, and I interact with 40S ribosomal subunit (*Chamond et al 2014*), and domain J-K recruits eIF4G, enhanced by eIF4A (*Pestova et al 1996; Lomakin et al 2000*). The structure of domain J-K-eIF4G1(HEAT1)-eIF4A revealed that positively charged patches on the eIF4G1-HEAT1 domain interact with two separated negatively charged clefts (in domains J and St) without perturbing its innate function of recruiting eIF4A (*Imai et al 2016; Imai et al 2023*). The binding sites for PTB1 were revealed to be dispersed, encompassing domains H to L (*Kafasla et al 2009*). These domains function as a single entity to form 48S PIC and any mutation in conserved IRES motifs can drastically affect the translation rates (*Fernández-Miragall et al 2009; Fernández et al 2011*). For example, the RAAA and GNRA loop in the domain I of type 2 IRES are crucial for IRES activity (*Robertson et al 1999*), and the GNRA loop is also found in other IRES families such as type 1 (poliovirus) and type 5 (Aichi virus A) picornaviruses (*Abdullah et al 2023*). The inherent flexibility within the IRES domains provides a challenge for structural studies of full-length IRES and to capture IRES in the context of 48S PIC. EMCV IRES can independently interact with 40S subunit without any eIFs (*Chamond et al 2014*) and does not require scanning to recognize the start codon (*Pestova et al 1996*), unlike type 1 IRESs, which scan for the start codon (*Sweeney et al 2014*). Besides, EMCV IRES can form 48S PIC with HEAT1-eIF4G without the requirement of eIF4G residues 1015-1104 that are known to interact with eIF3 (*Lomakin et al 2000*). These residues are indispensable in case of canonical initiation on capped mRNAs and for type 1 IRESs (*Villa et al 2013; Sweeney et al 2014*). However, how EMCV IRES interacts with 40S subunit, or what molecular interactions are essential to form the EMCV IRES-48S PIC remains a question.

In this study, we have used pull-down assay to isolate EMCV IRES-bound 48S PIC from Rabbit Reticulocyte Lysate (RRL) and subjected the complex to cryo-electron microscopy (cryo-EM) analysis. The cryo-EM map of EMCV IRES-bound 48S PIC, henceforth mentioned as EMCV IRES-48S PIC, shows densities corresponding to EMCV IRES domains that reveal how EMCV IRES contacts the ternary complex and 40S head at the inter-subunit interface. The structural details presented here account for numerous biochemical studies that have been reported earlier on Type 2 IRES. Furthermore, the structural analysis suggests how Type 2 IRES would interact with 43S PIC to form 48S PIC. Importantly, the study reveals a unique strategy used by viral IRES to capture the host translational apparatus for making viral polypeptide.

## Results

### 1. Overview and features of EMCV IRES-48S PIC

To isolate 48S PIC on EMCV IRES, we used a Talon affinity-based pull-down from nuclease-treated RRL. EMCV IRES RNA used harboured residues from 280 to 905 with AUG (start codon) at 834^th^ position. PTB1 was recombinantly overexpressed with an N-terminal 6X His tag, followed by a 3C protease cleavage site. PTB1 was incubated with the IRES, followed by RRL addition, and 48S PIC was stalled using GMP-PnP. The complex was eluted from the Talon matrix employing 3C protease cleavage and pelleted (Sup. Fig. 1.1 A). The EMCV IRES-48S PIC was subsequently analysed by Cryo-EM. The processed data yielded 3 major classes-(i) 40S without any factors-Map A, (ii) 40S-IRES-tRNA_i_-Map B, and (iii) 40S-IRES-ternary complex-Map B1, namely, EMCV IRES-48S PIC (Sup. Fig. 1.1 B). Map B and Map B1 have an overall resolution of 4.6 Å and 5.0 Å, respectively (Sup. Fig. 1.1 B). The core of 40S is at around 4.0 Å, and the local resolution across the maps was largely in the range of 4.0-8 Å (Sup. Figs. 1.2 A and B). Only the extreme tip of beak of 40S in Map B1 (Sup. Fig. 1.2 A) and ends of IRES and eIF2γ are around 12 Å in Map B2 (Sup. Figs 1.2 B).

The cryo-EM densities in EMCV IRES-48S PIC correspond to 40S, tRNA_i_, eIF2α, eIF2γ, and RNA in the mRNA channel, along with an extra density connecting 40S ribosomal head to tRNA_i_ (Fig. 1 A-B). However, it lacks distinct density for PTB1, eIF4G, eIF4A, and eIF3, and hence these factors are not modelled. Since nuclease-treated RRL lacked endogenous RNA, the presence of density for mRNA in the channel indicates trapping of EMCV IRES in 48S PIC. Also, the density connecting 40S head to tRNA_i_ in Map B and B1 could be assigned to a double-stranded RNA structure found in EMCV IRES (Fig. 1 A-B).

**Figure 1.**
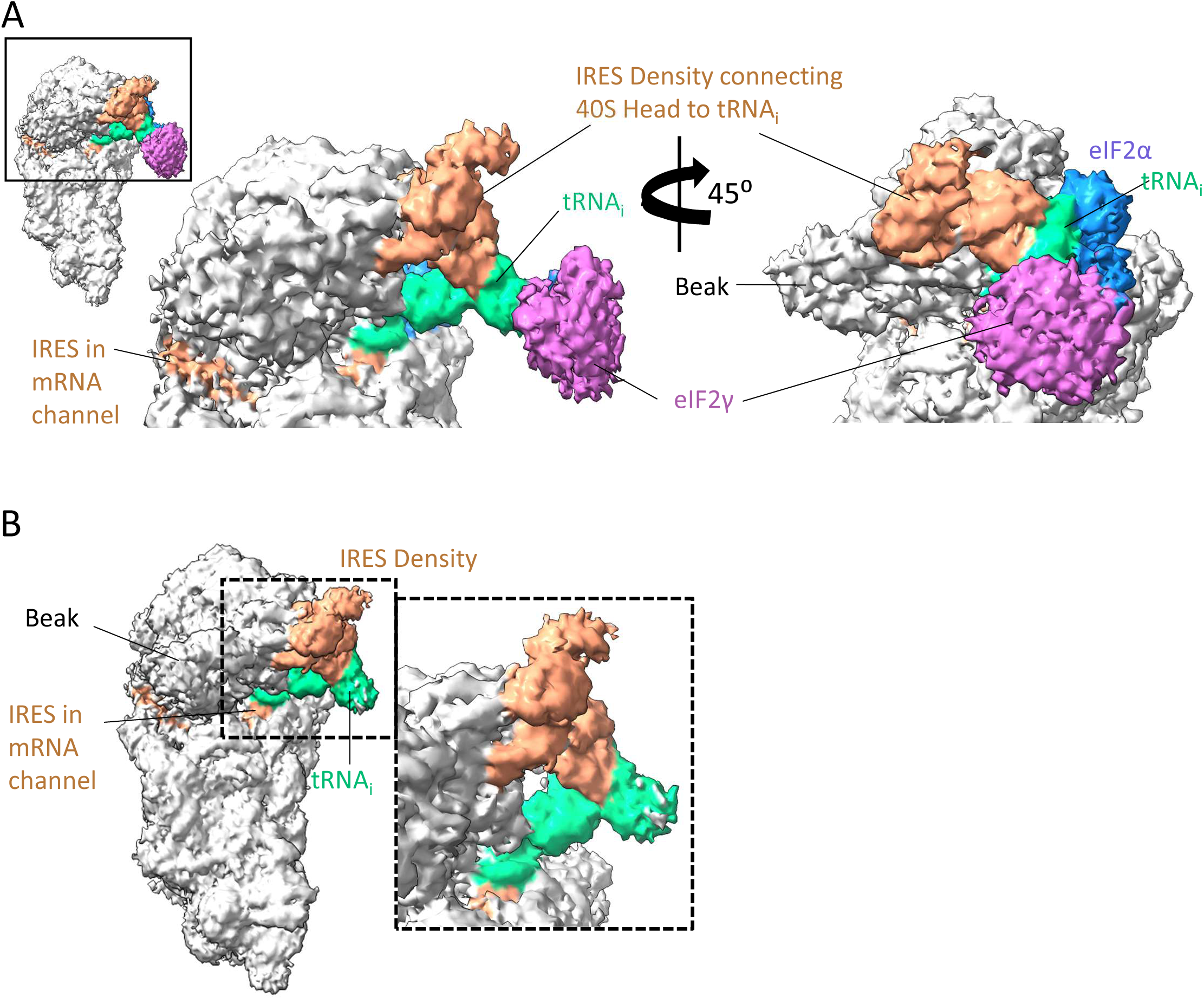
**(A)** Different views of EMCV IRES-48S PIC (Map B1) by 45⁰ rotation along one axis. The map shows densities assigned to 40S ribosome, RNA (in channel), ternary complex and IRES density at the inter-subunit region of head. **(B)** View of Map B and zoomed view of densities corresponding to IRES and tRNA_i_.

Besides, an extra density is evident at the mRNA entry site, contacting 40S ribosomal proteins-uS3, eS10, and uS5 and h16 (Helix 16) of 18S rRNA (Sup. Fig. 1.2 C). In the human 48S PIC, this region positions eIF3g-RRM (RNA recognition motif) and downstream ORF (Sup. Fig.1.2 D) (*Brito Querido et al 2020; Brito Querido et al 2024)*. We anticipate that this extra density in EMCV IRES-48S PIC could be contributed by downstream ORF; however, it’s difficult to assign it to eIF3g-RRM due to the lack of eIF3 (core/peripheral) density. Furthermore, our recent study on interaction of yeast eIF4B with 40S ribosome suggested occupancy of eIF4B (N terminal-RRM) in the same region (*Datey et al 2025*), which opens up the possibility of mammalian eIF4B-NT-RRM to bind to this region in the absence of eIF3, although in the human 48S PIC eIF4B was tentatively positioned slightly away from this location (*Brito Querido et al 2024)*. An interesting possibility could be positioning one of the RRM domains of PTB1 bound to UCUUU sequence of 18S rRNA (PTB1 can bind to UCUUU sequence-*Maris et al 2020*) present at the tip of h16 (528-532 nucleotides) of 18S rRNA. An extra density in the same position is also present in Map A (40S without EMCV IRES or factors-Sup. Fig. 1.2 E) as well, and PTB1 was used as a bait for the pull-down; therefore, this density may be contributed by PTB1-RRM interacting with h16 of 18S rRNA.

### 2. The EMCV IRES-48S PIC is trapped in a closed conformation

In EMCV IRES, A-834-U-835-G-836 (AUG-834) forms the start codon *(Kaminski et al 1994; Hellen and Wimmer 1995; Pestova et al 1996).* Previous experiments based on toeprints of EMCV IRES-48S PIC assembly suggested A-826-U-827-G-828 (AUG-826) as the codon where 48S PIC can assemble, and AUG-834 as the start codon (*Pestova et al 1996*). Furthermore, the intensity of the toeprint at AUG-834 was much higher than at AUG-826 (*Pestova et al 1996; Sweeney et al 2014*). Also, AUG-834 is present in a Kozak context (CRCCaugG; R is a purine) *(Kozak 1989)*, where the -3 position is A-831 and +4 is G-837 (Sup. Fig. 2 D), whereas AUG-826 is present in a poor Kozak context. In this work, the EMCV IRES construct used has both AUG-826 and AUG-834. Based on the above-mentioned reports, poor and strong Kozak context of AUG-826 and AUG-834, respectively, and placement of AUG-834 at the P site in EMCV IRES-40S binary complex (*Chamond et al 2014*), we reason AUG-834 to base-pair with the anticodon in EMCV IRES-48S PIC (Fig. 2 A), and the flanking nucleotide residues were added as per the sequence (Sup. Fig. 2 D). The recognition of the start codon at the P-site by tRNA_i_ leads to accommodation of ternary complex, and 48S PIC adopts a P_IN_ state in contrast to P_OUT_ state observed during scanning *(Llacer et al 2015; Llacer et al 2021; Yi et al 2022; Petrychenko et al 2024)*. A distinct difference of ∼7.0 Å could be seen by comparing the tRNA_i_ position of EMCV IRES-48S PIC with that in the open state (Fig. 2 B), depicting a P_IN_ state of tRNA_i_. The entrapment of the start codon in the P-site also evokes closure of the mRNA latch formed by 18S rRNA helices-h34 of 40S head and h18 of body (*Hussain et al 2014; Llacer et al 2015; Hinnebusch 2017*). The h34 of 18S rRNA of EMCV IRES-48S PIC is shifted by 9 Å as compared to the human canonical open state 48S PIC (PDB Id-7QP6, *Yi et al 2022*). This conformation of 18S rRNA correlates well with that of closed 48S PIC (PDB Id-7QP7, *Yi et al 2022*), suggesting EMCV IRES-48S PIC was captured in a closed state (Fig. 1 C). This closed conformation locks the mRNA in the channel formed within 40S head and body. On the other hand, the conformation of 18S rRNA in Map A (40S ribosome without initiation factors) shows an open state (Sup. Fig. 2 E). The transition from open (PDB Id-7QP6) to closed states (PDB Id-7QP7) structures is also accompanied by an upward shift of N terminal domain of eS17 (connecting the head to body) by ∼10 Å (*Yi et al 2022*). The EMCV IRES-48S PIC structure correlates with the conformation of eS17 in the human 48S-closed PIC, that is, eS17 N-terminal helix associated with the ribosomal head shifts upward by ∼10 Å, keeping C-terminal domain position constant (Fig. 2 D; Sup. Fig. 2 F).

**Figure 2.**
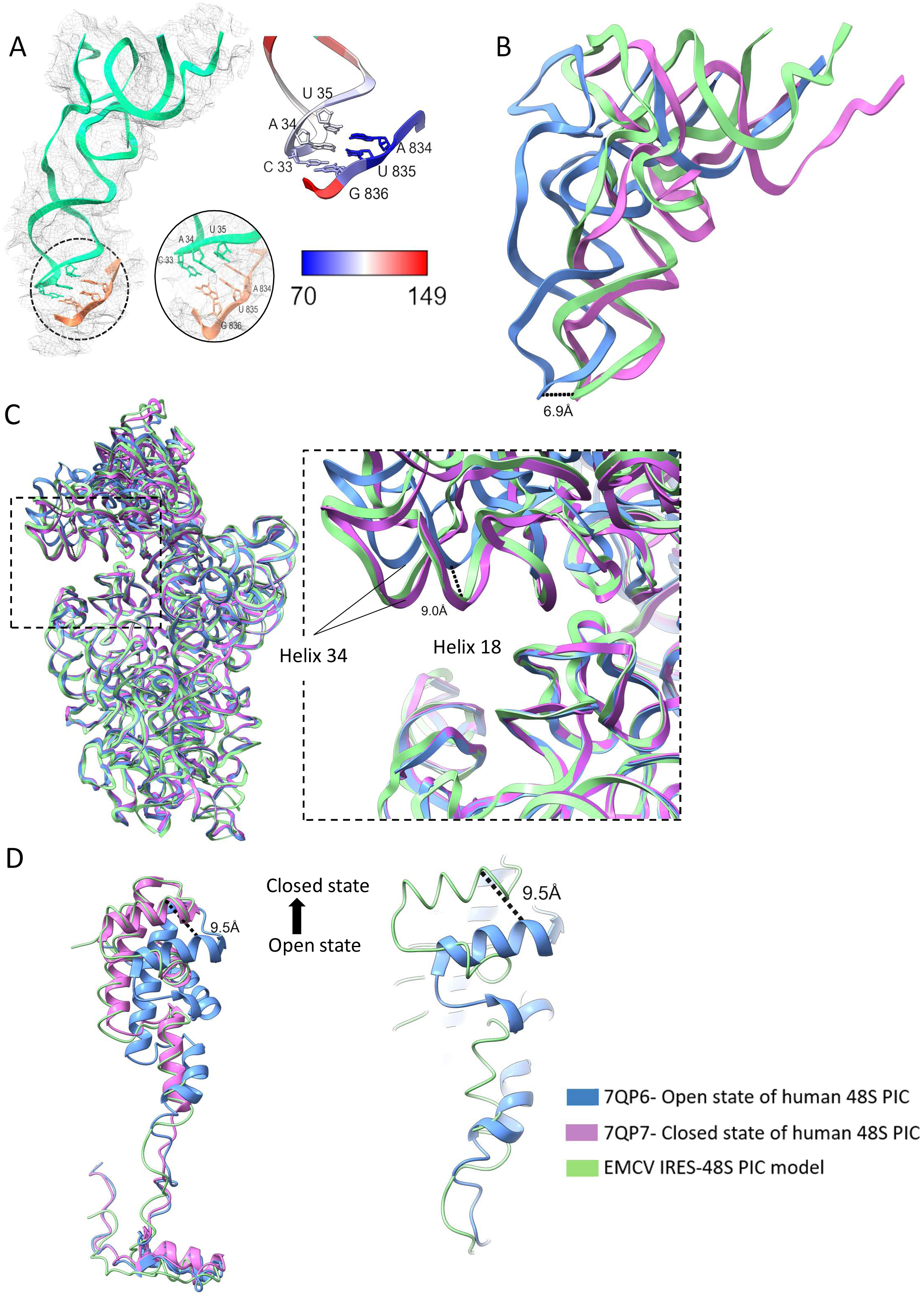
**(A)** Fitting of tRNA_i_-base paired to start codon AUG (left). Zoomed view of codon-anticodon interaction (middle), and B-factor for codon-anticodon interaction (right). **(B)** The tRNA_i_ in EMCV IRES-48S PIC is in P_IN_ state similar to that in human 48S PIC P_IN_ state (PDB Id-7QP7) and its anticodon shows ∼7 Å shift as compared to human 48S PIC open state (PDB Id-7QP6). **(C)** Comparison of ribosomal conformation (18S rRNA) of EMCV IRES 48S PIC with Human open and closed state (PDB Id-7QP6 and 7QP7, respectively). Focusing on the entry site and the helices governing latch conformation-Helix 34 moves toward helix 18 by 9 Å. **(D)** Movement of eS17 in open and closed states of ribosome. Zoomed view of comparison between eS17 in EMCV IRES 48S PIC and Human 48S PIC open state-showing an upward shift of the N-terminal domain by ∼10 Å.

### 3. The IRES density connecting 40S head to tRNA_i_ is contributed by the I domain apical part

In EMCV IRES-48S PIC complex, we observed a density, likely for double-stranded RNA, connecting 40S head to the tRNA_i_ elbow region (Fig. 3 A). The extra density interacts with the elbow region and acceptor stem of tRNA_i_ and ribosomal proteins-uS19 (RPS15) and uS13 (RPS18) (*Nomenclature-Ban et al 2014*). The EMCV IRES contains 4 major stem-loops (H-L) in the functional IRES region (Fig. 3 B) *(Hellen and Wimmer 1995)*. Among these, domains H and I have been shown to interact with 40S *(Chamond et al 2014)* and mutations of conserved residues in these domains severely compromise translation on EMCV IRES *(López de Quinto and Martínez-Salas 1997; Hellen and Wimmer 1995)*. Moreover, incubation of Foot and Mouth Disease Virus (FMDV) IRES, an EMCV-like type 2 IRES, with 40S ribosomes has shown a decrease in SHAPE reactivity in its domain 3 apex (*Lozano et al 2018*), which corresponds to EMCV IRES domain I apex. We reasoned that domains H and I may contribute to the double-stranded RNA density emanating from 40S head. The density architecture (Fig. 3 C) could be interpreted as a long main stem (S1) extending away from 40S where the base is anchored to the ribosome by two branches (B1 and B2), and to tRNA_i_ by one branch (B3), further divided into two sub-branches (B3a and B3b). Visualizing the IRES RNAfold determined-secondary structure (Fig. 3 D-which correlates with the experimental structure proposed in *Duke et al 1992*), this architecture could be contributed by the apical part of domain I.

**Figure 3.**
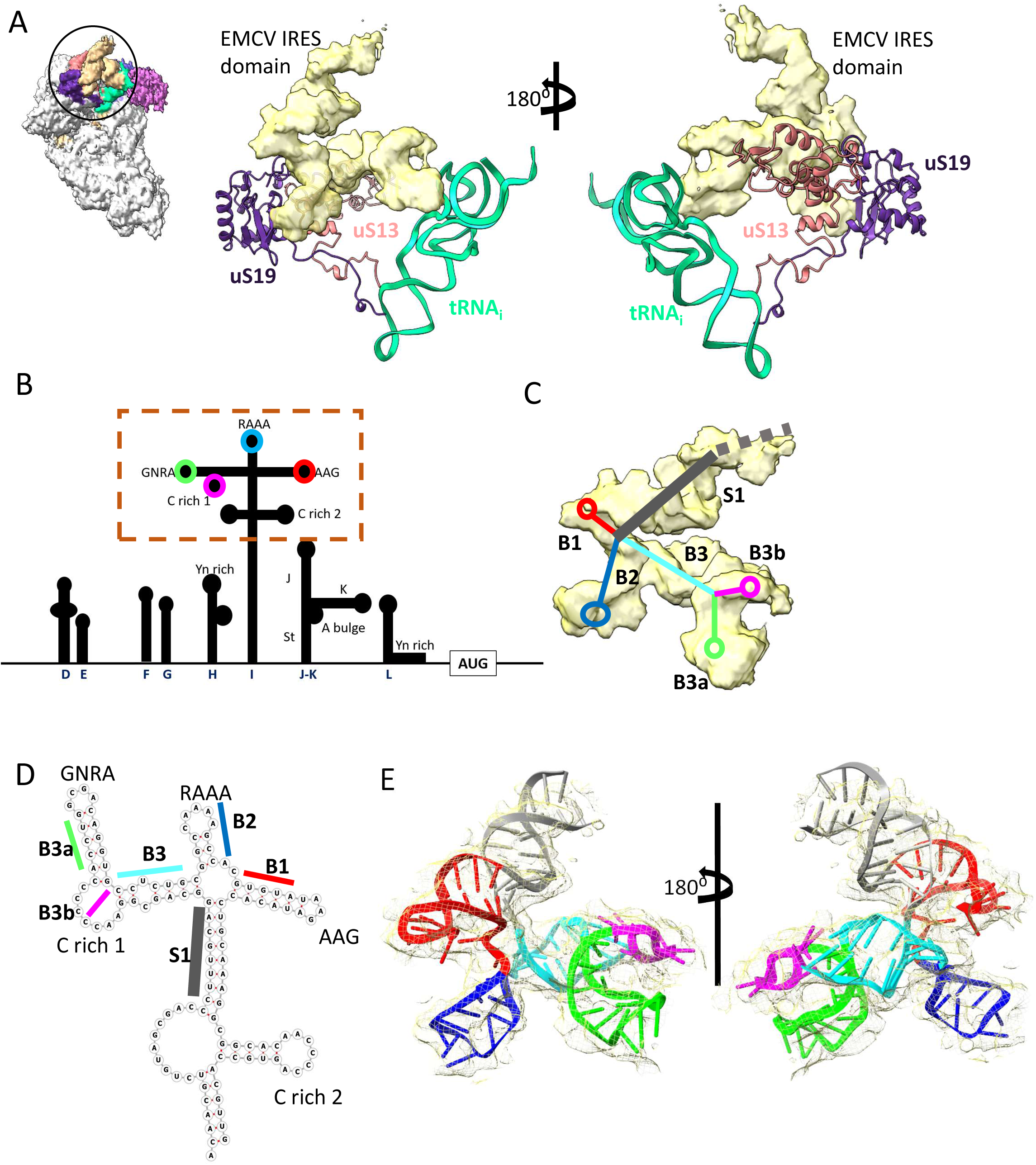
**(A)** Extra density connecting the head of 40S to tRNA_i_ elbow region is contributed by EMCV IRES RNA in Map B1. Rotated views of RNA density showing its connection to the 40S head via uS13 and uS19 and to tRNA_i_ via its elbow region. **(B)** Organization of EMCV IRES domains from D-L, where H-L makes the functional IRES moiety (*Hellen and Wimmer et al 1995*). **(C)** Deciphering the architecture of the obtained IRES density. The density could be interpreted as a long main stem (S1) extending away from the ribosome, where the base is anchored to the ribosome by two branches (B1 and B2), and to tRNA_i_ by one branch (B3), further divided into two sub-branches (B3a and B3b). **(D)** Secondary structure of apical region of domain I (made using RNAfold) marking the Stem and branches, along with imported loops. **(E)** Fitting of domain I apex in the density (Rotated views). The sub-domains are coloured as proposed in Fig 3C-D.

To determine the tertiary structure of the domain I apex, the IRES region from nucleotide 507-619 was modelled using Alphafold3 (Sup. Fig. 3.1 A). The model was decomposed and reconstructed based on the best-fit in the obtained density for the IRES using Chimera and Coot (Sup. Fig. 3.1 A). The final model was generated after multiple rounds of geometry correction and real space refinements (Fig. 3 E). The final model holds a correlation coefficient of 0.8 with respect to the map (Sup. Fig. 3.1 B), where B1 is AAG loop, B2 is CAAA loop, B3a-GCGA loop, B3b-C rich loop1, and S1 is the major double-stranded stem of domain I. The B-factor of the modelled IRES largely ranges from 124 to 200 (Sup. Fig. 3.1 C). To check the possibility of other IRES domains that might contribute to the extra density, Alphafold3 was used to predict the tertiary structure of isolated EMCV IRES domains (Sup. Fig. 3.2 A-J), using sequences as shown in Sup. Table 1. The predicted tertiary structure of domain H or experimental models of domains D to F did not fit in the observed IRES density (Sup. Fig. 3.2 K-L). The domain J-K adopts a Y-shaped structure, and placement of its cryo-EM (PDB Id-8HUJ) or NMR (PDB Id-2NBX) structure in the density would clash with 40S (Sup. Fig. 3.2 M). Moreover, in context to EMCV IRES-48S PIC, domain J-K binds eIF4G and the location of eIF4G has been mapped close to ES6 of 18S rRNA, located near the left foot 40S ribosome (*Yu et al 2011*). The domain I apex model in EMCV IRES-48S PIC shows the RAAA and AAG motif contacts uS19 and uS13, and the GNRA loop with tRNA_i_ (Fig. 4 A). In addition, incubation of EMCV IRES with rabbit reticulocyte lysate (RRL) protected domain I apex regions, including the CAAA loop in the SHAPE reactivity profile (*Maloney and Joseph, 2024*). These interactions with 40S head and tRNA_i_ could be facilitated by the long length and flexible nature of domain I.

**Figure 4.**
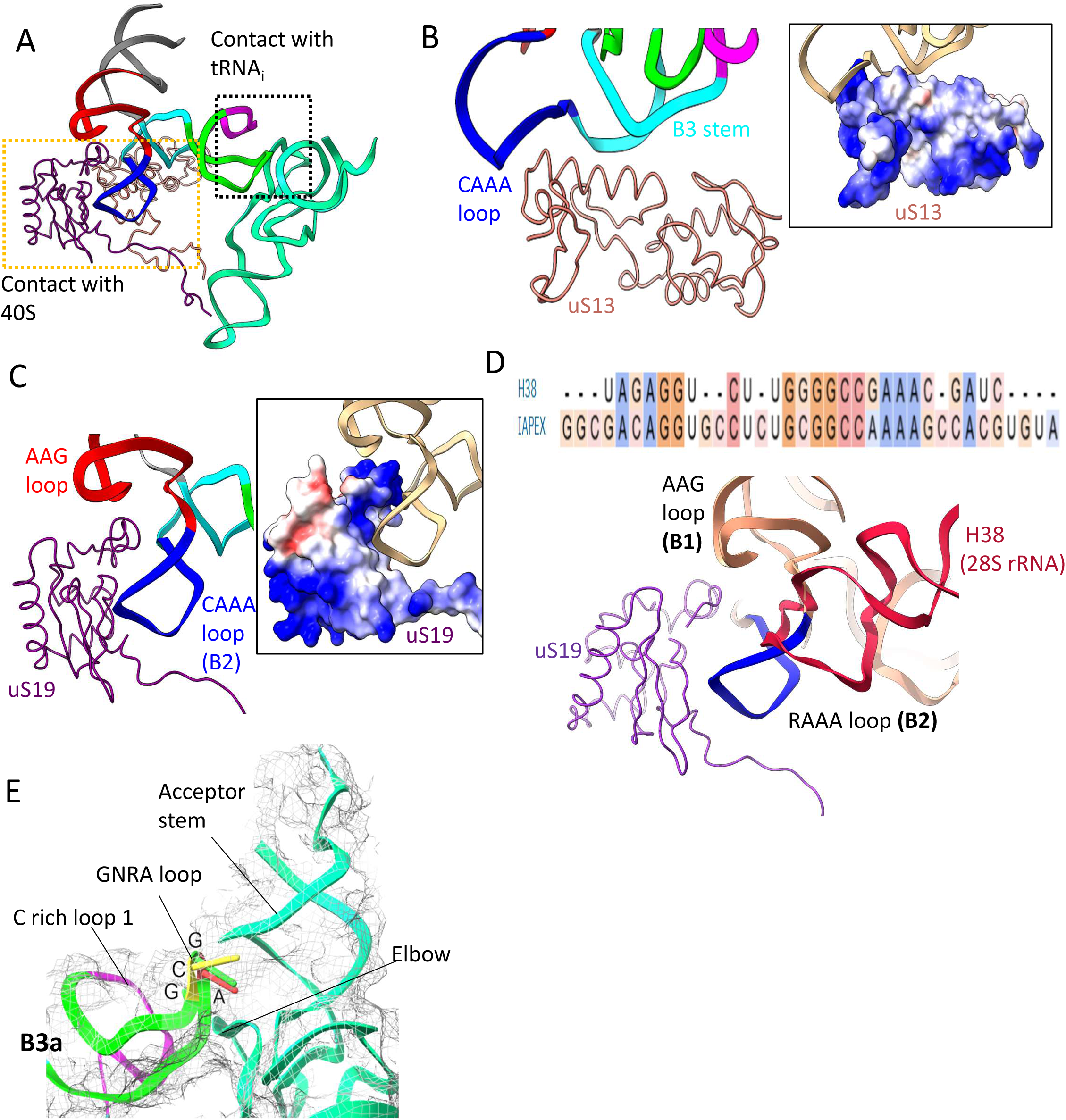
**(A)** Model showing connections of domain I apex with uS13, uS19, and tRNA_i_. **(B)** uS13 interacts with B3 branch or sub-domain of IRES via its alpha helix (100-117 residues). **(C)** Multiple point of contacts between uS19 and domain I motifs-RAAA and AAG. The electrostatic potential map of uS19 suggests that EMCV IRES interacts via ionic interactions with its phosphate backbone. **(D)** Sequence alignment of h38 with domain I apex of EMCV IRES showing sequence identity. Overlapping of uS19 from 80S (PDB Id-4UG0) and EMCV IRES-48S PIC show that the interaction of uS19 with EMCV IRES is similar to interaction with that of h38. **(E)** The fit of GNRA or GCGA stem and its contact with tRNA_i_ at the elbow region and acceptor stem.

### 4. The interaction of domain I of EMCV IRES with ribosomal proteins and initiator tRNA

The domain I is the longest domain in EMCV IRES, which harbours important motifs such as GNRA, RAAA, and C rich, crucial for IRES activity (*Roberts and Belsham, 1997; Fernández-Miragall and Martínez-Salas 2003*) and conserved across all cardioviruses (*Hellen and Wimmer 1995).* We mutated the GNRA loop and RAAA loop in EMCV IRES and checked for luciferase activity using a firefly *Luciferase* reporter downstream of the wild-type and mutant IRESs. We found a drastic reduction in the luciferase activity in the mutants as compared to that of the wild-type (Sup. Fig. 4 A), which correlates with previous studies that showed the importance of these motifs in regulating IRES activity (*Roberts and Belsham, 1997; López de Quinto and Martínez-Salas, 1997; Robertson et al 1999; Fernández-Miragall and Martínez-Salas 2003*). Among these, the CAAA and AAG motifs share potential contact sites with uS13 and uS19. The alpha helix (100-117 residues) of uS13 contacts the IRES element at the B3 stem (connecting the GNRA loop to RAAA loop) (Fig. 4 B). uS19 contacts the IRES majorly at its CAAA motif through multiple sites involving N-terminal residues, residues-67 to 75, and C-terminal-102 to 124 (Fig. 4 C). These regions of uS13 and uS19 are rich in basic residues, which might interact with the negatively charged backbone of the IRES element (Fig. 4 B-C). The role of uS13 and uS19 also involves the formation of inter-subunit bridges during 60S joining to form elongation-competent 80S complexes. uS13 interacts with uL5 (RPL11) and uS19 with helix 38 or h38 (1748-1778, in humans) in 28S rRNA to form inter-subunit bridges-B1b/c and B1a, respectively (Sup. Fig. 4 B) (*Ben-Shem et al 2011; Bowen et al 2015; Khatter et al 2015*). These interactions are dynamic owing to ribosomal subunit rotation and swivelling during 80S ribosomal translocation states (*Khatter et al 2015*).

On superimposition of 80S ribosomal structure to EMCV IRES-48S PIC model, the IRES density clashes with the position of uL5 and h38 of 28S rRNA (Sup. Fig. 4 C), suggesting repositioning of IRES domain from 40S head during 48S to 80S transition. Interestingly, the h38 residues interacting with uS19 shares considerable similarity in sequence to I domain apex in the EMCV IRES (Fig. 4D), suggesting the domain I apex of EMCV IRES could mimic h38 (60S)-40S interaction (Fig. 4D). The similarity of h38 with the domain I residues provides additional support for annotation of domain I apex in the density.

The GCGA (GNRA motif-where N is any nucleotide and R is a purine) is known for long-range RNA-RNA interactions, widespread in ribosomal RNA and in some catalytic RNAs. It forms a characteristic U-turn structure (*Fiore and Nesbitt 2013*) and interacts with minor grooves of helical RNA elements (Sup. Fig. 4 D-*Reiter et al 2010*). A single point mutation within this tetraloop (GCGA to GCGC) severely reduced the IRES activity, suggesting its essential for IRES activity (*Roberts and Belsham, 1997; Fernández-Miragall and Martínez-Salas 2003*). The density extending from the elbow region of tRNA_i_ could fit in the characteristic U-turn, adopted by conventional GNRA motifs. In EMCV IRES, the GNRA motif is represented by GCGA loop, preceded by a C-rich region and in close contact with the tRNA_i_ elbow and acceptor stem (Fig. 4 E). Thus, we infer that EMCV IRES interacts with tRNA_i_ by virtue of its GCGA loop.

### 5. The position of eIF2-ternary complex is shifted towards 40S head in EMCV IRES-48S PIC in contrast to canonical 48S PIC

We could fit eIF2α and eIF2γ in their respective densities in Map B1 (Fig. 5 A). Focussed classification or refinement did not yield any distinct density corresponding to the position of eIF2β, probably due to the flexibility associated with repositioning of eIF2β during transition from open to closed complexes *(Llacer et al 2015; Llacer et al 2021; Yi et al 2022; Petrychenko et al 2024)*. Previous reports on EMCV IRES suggested its direct interaction with eIF2 (*Scheper et al 1991; Scheper et al 1994*) and inactivation of eIF2 compromises EMCV IRES-mediated translation *(Welnowska et al 2011; Kwon et al 2016)*, indicating EMCV IRES’s dependence on the canonical ternary complex. Here, we observe a direct interaction of EMCV IRES with ternary complex via tRNA_i_, a feature not observed in previously determined HCV (*Yamamoto et al 2014; Yamamoto et al 2015; Brown et al 2022*) and CrPV IRES-bound ribosomal structures (*Pisareva et al 2018; Acosta-Reyes et al 2019*).

**Figure 5.**
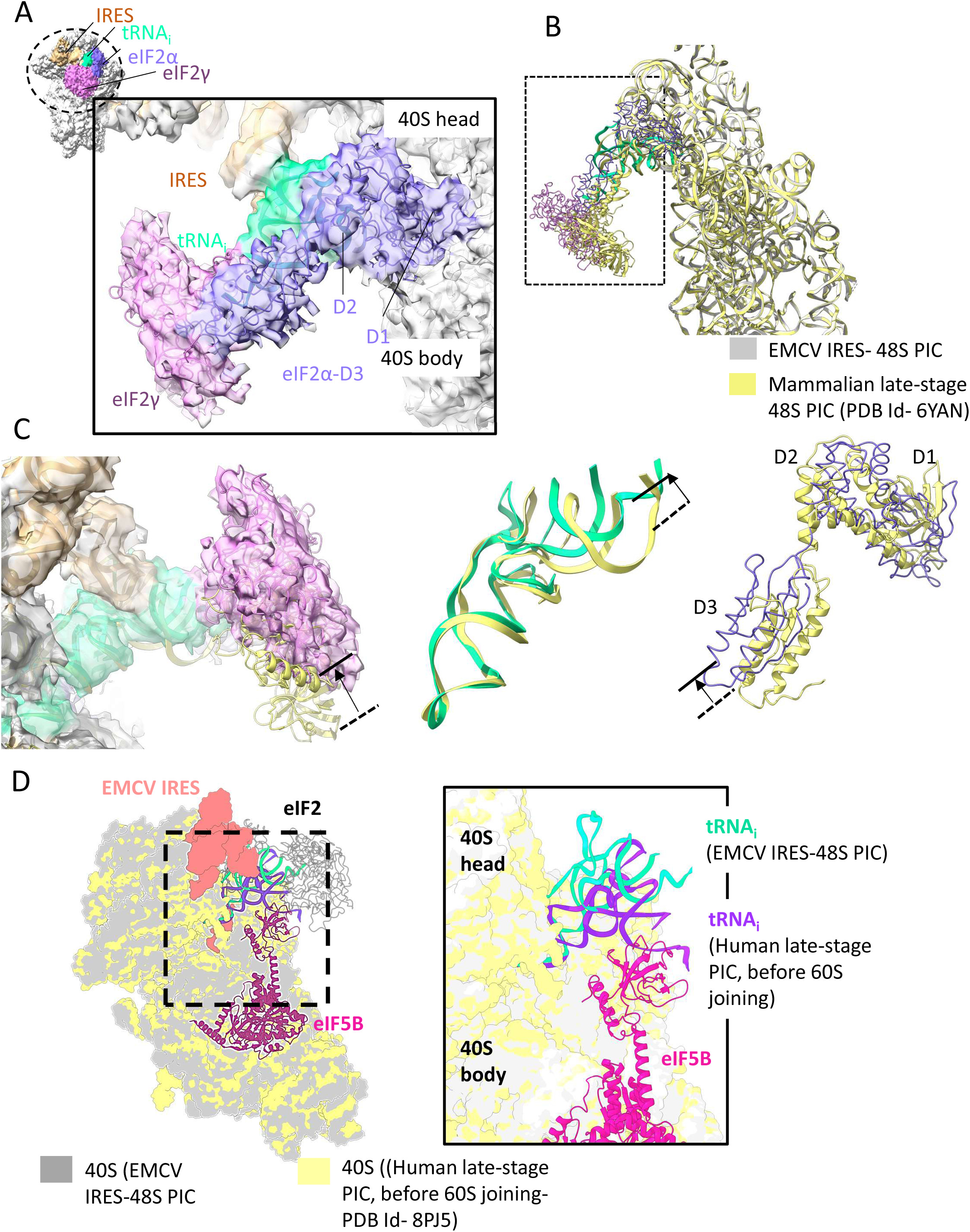
**(A)** Inter-subunit view of EMCV IRES 48S PIC showing position for ternary complex on map. Fitting of eIF2α and eIF2γ in its corresponding density in Map B1. **(B)** Overlapping EMCV IRES 48S PIC on mammalian late-stage 48S PIC (PDB Id-6YAN), indicates a shift in position of eIF2γ towards 40S head. **(C)** Zoomed view showing the positions of eIF2γ, tRNAi, and eIF2α in the EMCV IRES 48S PIC relative to those in the mammalian late-stage 48S PIC (PDB ID 6YAN). The black arrow indicates the shift in position. **(D)** Superimposition of the EMCV IRES-48S PIC on the human late-stage 48S PIC (before 60S joining; PDB Id-8PJ5), showing the conformation of tRNA_i_ in association with eIF2 and IRES and tRNA_i_ with eIF5B in the canonical context, respectively. (right) Zoomed view of tRNA_i_ conformation in both complexes.

As found in canonical 48S PICs (*Hashem et al 2013*; *Brito Querido et al 2024; Petrychenko et al 2024)*, eIF2α-domain 1 is in close contact with ribosomal protein-uS7, and domain 2 with tRNA_i_ elbow and domain 3 with eIF2γ in EMCV IRES-48S PIC. The position of eIF2γ and eIF2α-domain 3 (D3) is distinct from mammalian 48S-closed PICs (PDB Id-6YAN and 7QP7) as we could observe a shift by ∼10 Å towards the head of 40S on superimposing the 18S rRNA from EMCV IRES 48S PIC and mammalian late-stage 48S PIC (Fig. 5 B). The ternary complex is flexible, and it moves away from ribosomal head towards the body on recognition of authentic start codon as studied in yeast (*Villamayor-Belinchón et al 2024*) and human (*Petrychenko et al 2024)*. This opposite directional shift of the ternary complex in the EMCV IRES-48S PIC is evident in tRNA_i_ acceptor stem as well (Fig. 5 C). This shift could be due to the rigid stem B3 (consisting G-C base pairs), connecting 40S head to GNRA loop, which interacts with the tRNA_i_ at its elbow and acceptor arm, and the association of eIF2α-D3 and eIF2γ with the acceptor arm of tRNA_i_ orchestrated with the conformational change.

During the transition of 48S PIC to 80S elongation-competent complex, there are major changes in the conformation of tRNA_i_ due to the joining of eIF5B, and release of eIF2 (*Petrychenko et al 2024*). This joining event of eIF5B positions the tRNA_i_ elbow and acceptor stem towards the 40S body to aid 60S ribosomal subunit joining (*Petrychenko et al 2024*). However, in the context of EMCV IRES-48S PIC, we have seen the positioning of tRNA_i_ elbow and acceptor stem towards the 40S head, away from the body (Fig. 5 C). On superimposing the human 48S PIC structure (before 60S joining), 48S-5 (PDB Id-8PJ5-*Petrychenko et al 2024*), we could see that tRNA_i_ in EMCV IRES-48S PIC is away from the canonical tRNA_i_ position (in contact with eIF5B) (Fig. 5 D). Therefore, we anticipate a change in tRNA_i_ conformation during eIF5B joining and eIF2 release. Furthermore, the IRES (domain I) interacting with the tRNA_i_ elbow needs to be displaced from the position to facilitate the interaction of tRNA_i_ with eIF5Band this rearrangement would also aid in 60S joining and avoid steric clash with the IRES domain I.

## Discussion

The mechanism of IRES recruitment on 40S ribosome varies considerably among different types of IRESs. CrPV intergenic IRES binds to 40S ribosome by mimicking tRNA-mRNA interaction with the help of pseudoknot 1 *(Petrov et al 2016; Fernandez et al 2014; Murray et al 2016; Acosta-Reyes et al 2019)*, whereas HCV IRESs associates with the solvent side of 40S body by replacing eIF3 with its domain 3 *(Spahn et al. 2004; Hashem et al 2013; Niepmann and Gerresheim 2020; Brown et al 2022)*. In this study, we capture EMCV IRES in 48S PIC context, interacting with 40S ribosomal head and tRNA_i_. Model building suggests that domain I of EMCV IRES interacts with 40S ribosome head and tRNA_i_ elbow stem (Fig. 6 A). The similarity in the sequence of h38 of 28S rRNA with the domain I and its ability to interact with uS19-N terminal via RAAA motif (*Khatter et al 2015*) suggests a mimicry mechanism adopted by EMCV IRES for its recruitment to the ribosome. Moreover, the conservation of domain I apex sequence and motifs (RAAA, AAG loop, C-rich and GNRA) across Cardioviruses-Theiler’s murine Encephalitis virus (TMEV), Vilyuisk human encephalomyelitis virus (VHEV), Theiler’s-like rat virus (TRV) and Saffold viruses 1 and 2 (SAFV-1 and SAFV-2) and Apthoviruses (AAG loop is replaced by ACG loop)-Foot and mouth disease virus (*Hellen and Wimmer 1995; Liang et al 2008*) suggests that these IRESs might adopt similar strategies for 48S formation (Fig. 6 B-C). Like EMCV IRES, the type 1 IRES (Poliovirus, Coxsackie virus, etc.) also harbours the GNRA loop, preceded by a C-rich loop at its longest domain, known for long-range RNA-RNA interactions. The segment harbouring GNRA loop is highly conserved across the type 1 family of IRESs (*Kim et al 2015*). The domain I of EMCV IRES is similar to domain IV of polioviral IRES or other type 1 IRESs in terms of length, secondary structure, and conserved motifs (GNRA, C-rich) positioning (Fig. 6 C). Therefore, we anticipate a similar interaction of domain IV (in type 1 IRES class) with tRNA_i_. Also, this interaction of IRES with tRNA_i_ could be a strategy by which these IRESs can sequester the tRNA_i_ pool in the cell, rendering them unavailable for capped cellular mRNAs. During the revision of this work, a preprint reported a structure of polioviral IRES-48S PIC (*Velazquez et al 2025*), which shows that domain IV apex (similar to domain I apex in EMCV IRES) interacts with uS13 and uS19, and the GNRA loop directly interacts with tRNA_i_ during start codon recognition (*Velazquez et al 2025*) as observed in EMCV IRES-48S PIC. Similarly, the Aichi viral IRES (type 5-*Abdullah et al 2023*) harbours a GNRA loop in its longest domain, which is domain J. Deletion of the GNRA loop compromises the IRES activity; however, substitution mutations in this region either elevate the IRES activity or it remains unaltered (*Yu et al 2011*). We hypothesize that Aichiviral IRES might use this motif to mediate long-range interactions with tRNAi, similar to type 1 and type 2 IRESs, as all these IRESs require eIF2-ternary complex for the formation of 48S PIC.

**Figure 6.**
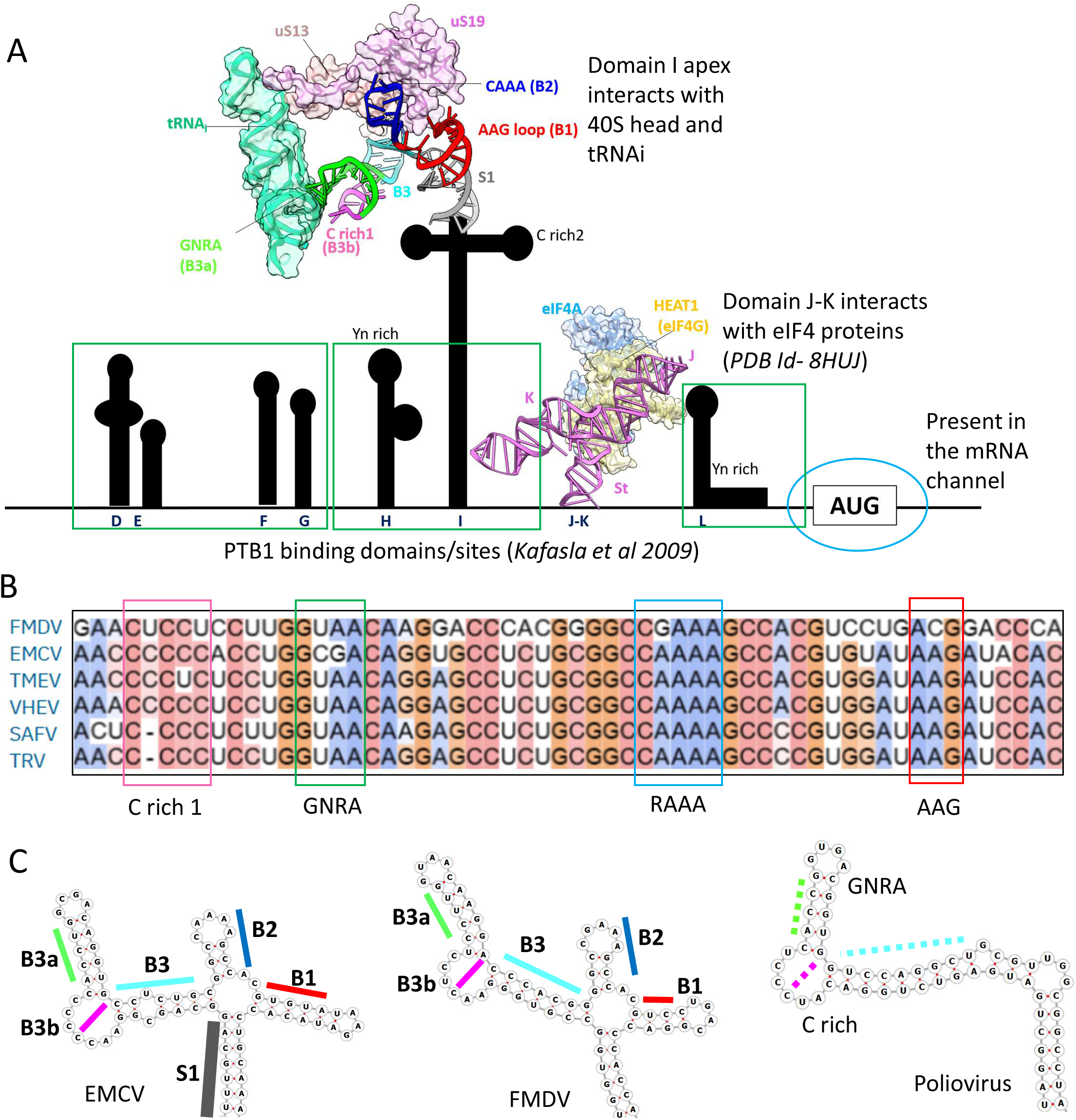
**(A)** EMCV IRES secondary structure depicting the role of binding partners for each domain in context to translation initiation. The known tertiary structure of EMCV IRES and its binding partners are depicted. The domains responsible for binding PTB1 are boxed. **(B)** Conservation of domain I apex sequence and secondary structure across type 2 IRES family. **(C)** Comparison of secondary structure of EMCV and FMDV IRES to that of Polioviral IRES (type 1).

The EMCV IRES does not require scanning, and the start codon (A-834) is directly placed in the P site, which would eventually place the domain L at the mRNA exit site, preceded by domain J-K that interacts with eIF4G-eIF4A (Fig. 6 A). Earlier, biochemical studies suggested eIF4G to be positioned close to ES6 of 18S rRNA in EMCV IRES-bound 48S PIC (*Yu et al 2011*). The human 48S PIC with a 5’ capped mRNA showed a similar location for eIF4G, that is at the mRNA exit site contacting eIF3 (*Brito Querido et al 2020; Brito Querido et al 2024)*. Locating eIF4F has been challenging due to the inherent flexibility associated with the eIF4F complex on mRNA and requires association with eIF3 in canonical 48S context (*Brito Querido et al 2020; Brito Querido et al 2024)*. However, the canonical eIF3-eIF4G interaction (Villa et al 2013) is dispensable for EMCV IRES-48S PIC formation (Lomakin et al 2000; Sweeney et al 2014), and no density for eIF3 was observed even after focused classification. However, after the initial submission of this work, a preprint reported a structure of reconstituted EMCV IRES-48S PIC where eIF3 is present at the canonical position (*Bhattarcharjee et al 2025*). This position of eIF3 suggests the possibility that eIF4G-eIF4A proteins could be placed similarly to the canonical eIF3-eIF4G-eIF4A position (*Brito Querido et al 2024*) in context to EMCV IRES-48S PIC. Thus, placing eIF4G-domain J-K close to ES6 of 40S ribosome, which coincides with the previous hydroxyl radical cleavage assay (*Yu et al 2011*). In addition to initiation factors, the ITAF-PTB1 serves as an essential ITAF for 48S PIC formation on EMCV IRES, but the obtained map shows no distinct density to PTB1. PTB1 binds to the base of domains H and I, and domain K loop (*Kafasla et al 2009; Dorn et al 2023*), and the flexibility associated with these domains might have hindered capturing of PTB1 in the reported 48S complex (Fig. 6 A). However, the unassigned extra density at the mRNA entry site could be contributed by PTB1-RRM interacting with 18S rRNA, as discussed previously.

The structural studies on type 2 IRESs have been limited due to their flexible nature. In the cryo-EM map and structural analysis presented here, we could capture a part of EMCV IRES in 48S context (Fig. 6 A) and fetch a significant understanding of ribosomal recruitment by EMCV IRES and 48S PIC formation. As mentioned above, the cryo-EM map of reconstituted EMCV IRES-48S PIC (*Bhattarcharjee et al 2025*) show similar findings, where the density for the apical portion of domain I of the IRES is only observed and it interacts with uS13 and uS19 on 40S ribosome, and tRNA_i_. A higher resolution of the reconstituted EMCV IRES-48S PIC (3.2 Å) helped the authors to identify interactions of individual IRES nucleotides with their binding partners (*Bhattarcharjee et al 2025*). In future, the entrapment of additional factors would significantly provide more insights into EMCV IRES 48S PIC. The conservation of secondary structures and motifs such as GNRA within the picornaviruses (type 1, 2, and Aichiviral IRESs) suggests common strategies of interaction in the context of 48S PICs in the Picornaviridae family.

## Materials and Methods

### 1. Plasmid constructs and Molecular Cloning

EMCV IRES 905 was obtained from EMCV-L plasmid-EMCV IRES (nt 280–905) into pCR2.1 and inserted in pcDNA3.1 using BamHI and XbaI restriction sites. Polypyrimidine tract binding protein 1 was cloned from HEK293 cDNA, and 3C protease site (LEVLFQGP) was inserted at the N-terminal and inserted into pET28a in between BamHI and HindIII restriction sites, retaining the N-terminal 6X Histidine tag.

EMCV IRES-Luciferase constructs-The firefly luciferase gene was inserted downstream of EMCV IRES in-frame with A-834 residue of IRES to generate the wild-type EMCV IRES-Luciferase construct in pcDNA3.1 (WT-Luc). The CAAAA (RAAA) loop and GCGA (GNRA) loop were mutated to GCTGA and TACG, as per functional assay reports for FMDV IRES and EMCV IRES to generate RAAAmut-Luc and GNRAmut-Luc, respectively (*López de Quinto and Martínez-Salas, 1997; Fernández-Miragall and Martínez-Salas, 2003*). The sequence of oligos or primers used is listed in Sup. Table 2.

### 2. Protein overexpression and purification

PTB1 (Histidine tag-3Cprotease site-*PTB1* gene) was overexpressed using 0.5 mM IPTG in *E. coli* BL21 cells at 30 ⁰C, 120 rpm for 4 hours in Luria broth. 2 l harvested culture was lysed using Buffer N1 (20 mM HEPES pH-7.4, 300 mM KCl, 2 mM MgCl_2_, 10 % glycerol, 10 mM imidazole, 5 mM β-mercaptoethanol, 0.05 % Triton-X-100, 2 mM PMSF) and sonicated at 18% amplitude, 10 s ON and 20 s OFF pulses, 30 cycles and centrifuges at 20000 rpm for 20 minutes and the supernatant was loaded on Ni-NTA column and eluted using a gradient of N1 to N2 buffer (20 mM HEPES pH-7.4, 300 mM KCl, 2 mM MgCl_2_, 10 % glycerol, 500 mM imidazole, 5 mM β-mercaptoethanol). PTB1 was eluted at 250 mM Imidazole concentration. The eluant fractions (diluted to 100 mM KCl) were further loaded on Heparin column and eluted by applying a gradient of 100 mM to 1000 mM KCl. The protein was eluted at 250 mM KCl concentration. The eluant fractions were stored at -80 ⁰C after Size Exclusion Chromatography in Buffer S (20 mM HEPES pH-7.4, 200 mM KCl, 2 mM MgCl_2_, 5 % glycerol, 1 mM DTT) at 2.5 mg/ml.

3C protease was overexpressed using 0.5 mM IPTG in *E. coli* BL21 cells at 30 ⁰C, 120 rpm in Luria broth. 2 l harvested culture was lysed using Buffer N11 (20 mM HEPES pH-7.4, 200 mM KCl, 2 mM MgCl_2_, 10 % glycerol, 10 mM imidazole, 5 mM β-mercaptoethanol, 0.05 % Triton-X-100, 2 mM PMSF) and sonicated at 18 % amplitude, 10 s ON and 20 s OFF pulses, 30 cycles and centrifuges at 20000 rpm for 20 minutes and the supernatant was loaded on Ni-NTA column and eluted using a gradient of N11 to N22 buffer (20 mM HEPES pH-7.4, 200 mM KCl, 2 mM MgCl_2_, 10 % glycerol, 500 mM imidazole, 5 mM β-mercaptoethanol). 3C protease was eluted at 250 mM imidazole concentration. The eluant fractions were subjected to Size Exclusion Chromatography in Buffer S (20 mM HEPES pH-7.4, 100 mM KOAc, 2 mM MgCl_2_, 10% glycerol, 1 mM DTT), and peak fractions were concentrated and stored at 2.5 mg/ml.

### 3. *In vitro* transcription of EMCV IRES

EMCV IRES 905-pcDNA3.1 was linearized using Xba1 restriction enzyme and transcribed using Promega-RiboMAX™ Large Scale RNA Production Systems as per manufacturer’s protocol. 1 µg of linearized plasmid yielded 87 µg of RNA after DNase treatment and RNA cleanup (RNA clean up kit-NEB). WT-Luc, GNRAmut-Luc, and RAAAmut-Luc were linearized using XhoI and transcribed *in vitro* using the same strategy.

### 4. Assembly of EMCV IRES 48S PIC using Talon Affinity Chromatography

12 µg of IRES was heated at 95 ⁰C to dissolve any secondary structures acquired and refolded using buffer R (20 mM HEPES pH-7.4, 150 mM KOAc, 2 mM MgCl_2_, 2 mM β-mercaptoethanol, 0.25 mM spermidine) at 37 ⁰C for 5 minutes, and PTB1 (1:2 = IRES: PTB1) was added and further incubated at 30⁰C for 5 minutes. Simultaneously, Rabbit Reticulocyte Lysate (RRL) (Promega) was incubated at 30⁰C with ATP, Amino acid mix minus Leucine, murine RNase Inhibitor for 5 minutes, and then mixed with the IRES-PTB1 vial with instant addition of 6 mM GMP-PnP per reaction (100 µl) and incubated at 30 ⁰C for 8 minutes, followed by ice incubation. The reaction was loaded onto Talon beads equilibrated with Buffer A (20 mM HEPES, pH 7.4, 150 mM KOAc, 2 mM MgCl2, 4% glycerol, 5 mM imidazole, 2 mM β-mercaptoethanol). After recommended passes, reloading and incubation, the flow through was collected and the beads were washed with Buffer A until A260 attains a baseline value (∼0), following which 3C protease in Buffer A was added to the beads and left for overnight incubation. Fractions were eluted using Buffer A (400 µl each in 10 vials) and subjected to analysis such as A260 measurements and agarose gel electrophoresis. Samples having the RNA bands were pelleted using a 1ml sucrose cushion (20 mM HEPES pH-7.4, 150 mM KOAc, 2mM MgCl_2_, 30% Sucrose, 1mM DTT) in SW60 tubes and centrifuged at 50,000 rpm for 10 hours at 4 ⁰C. The pellet was resuspended in 20 µl buffer R and used for Cryo-EM grid preparation (no crosslinker was used to avoid artifacts).

### 5. Negative stain analysis and Mass Spectrometry

The final sample was diluted 10 times using buffer R and applied on 400-mesh Cu TEM grids, which were freshly glow-discharged (negative polarity) for 30 s in GloQube glow-discharge system, stained using 1% Uranyl acetate solution, and analysed using Talos L120C transmission electron microscope at 57000 X magnification. Furthermore, the sample was digested by trypsin and subjected to NanoOrbitrap analysis for the identification of proteins in the complex.

### 6. Cryo-EM sample preparation

3 µl of resuspended pellet (2.96 A260) was applied on glow-discharged Quantifoil R 1.2/1.3 300 mesh 2nm Carbon Coated Grid and Blotted using 8s and 8.5s Blot time, zero Blot force at 16⁰C and 100% Humidity, and plunged into liquid ethane. Cryo-EM data were collected on Talos Arctica transmission electron microscope equipped with a FEG at 200 kV (Thermo Fisher Scientific). All data were collected using a Gatan K2 Summit Direct Detector at a nominal magnification of 36,000 X, and a pixel size of 1.17 Å with a total electron dose of 55 e-/Å2 fractionated over 20 frame movies with a dose rate of ∼2.5 e-/Å^2^ /frame.

### 6. Data Processing

Micrographs were collected and processed using CryoSPARC v3.3 (*Punjani et al 2017*). The micrographs were patch motion corrected and CTF was estimated. Using a blob picker with an average diameter of 300 Å particles were picked and extracted. The extracted particles were subjected to multiple rounds of 2D classification. Final 2D classes showing promising ribosome 2D features were selected for *ab initio* reconstruction. The junk was discarded, and the 237054 good particles were classified into 2 classes using a mask around tRNA_i_. We obtained 2 classes-Empty 40S (1.25L particles-Class 1 or Map A), and 40S with tRNA bound (1.11L particles-Class 2). Class 2 was further classified into 2 classes using a 3D mask around the IRES density and subjected to non-uniform refinement with global CTF refinement to yield Map B, having tRNA_i_ and IRES density (55k particles). This was subjected to Non-uniform Refinement (*Punjani et al 2020*) with global CTF refinement. Map B class was further classified using a mask around eIF2α and eIF2γ to yield 2 classes-40S-tRNA_i_-IRES-eIF2 (28k particles-Map B1) and 40S-tRNA_i_-IRES (26k particles). The obtained maps were then subjected to model building and refinement. The local resolution for the obtained maps were estimated using Phenix (*Liebschner et al 2019)*.

### 7. Map analysis and Model building

To the obtained maps, 40S ribosome (PDB Id-6YAN) was fitted using UCSF Chimera. The 40S head (Head proteins + 18S rRNA (1197-1688) + eS17) and 40S body (Body proteins + 18S rRNA (1-1196; 1689-1870)) was fitted to the obtained maps separately and subjected to Rigid Body Fit and Real Space Refinement using PHENIX *(Liebschner et al 2019)*. The models were merged using Coot *(Emsley et al 2010)* and subjected to Real Space Refinement. tRNA_i_ and mRNA models were taken from PDB Id-8OZ0 and fitted to the maps-B and B1 in Chimera. eIF2α and eIF2γ from PDB Id-8OZ0 were rigid-body fitted to Map B1 and then mutated as per Rabbit eIF2 protein sequence (NCBI Reference Sequence-XP_002719561.1; XP_051683593.1) and subjected to a final Real Space Refinement. The IRES I domain apex model was predicted from Alphafold3 (*Abramson et al 2024*) and the helical sub-domains were dismantled and fitted according to best fit using Chimera, followed by chain joining in Coot with manual Real Space Refinement (*Afonine et al 2018*). The geometry was corrected using Geometry minimization tool in Phenix with rounds of Real Space Refinement. H domain model was predicted using Alphafold3. The final model yielded 40S-EMCV IRES-tRNA_i_ (Map B) and 40S-EMCV IRES-tRNA_i_-eIF2αϒ (Map B1). All the figures were made using ChimeraX *(Pettersen et al 2021).* The models were real space refined using Phenix, and the Fourier Shell Correlation for ‘Map to Model’ for each was determined at 0.5 FSC (Sup. Fig. 2A-C; Table 1).

**Table 1:**
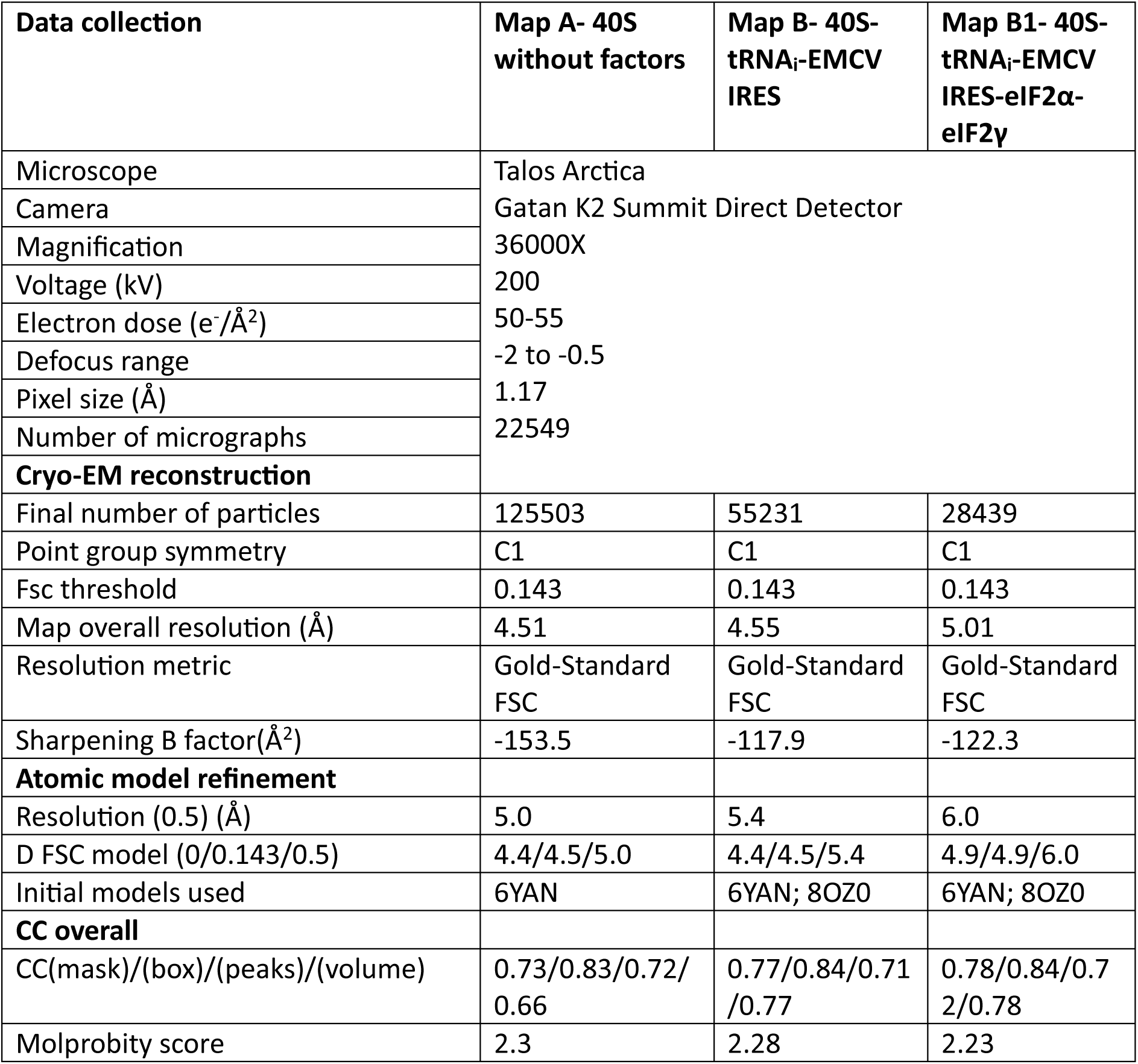
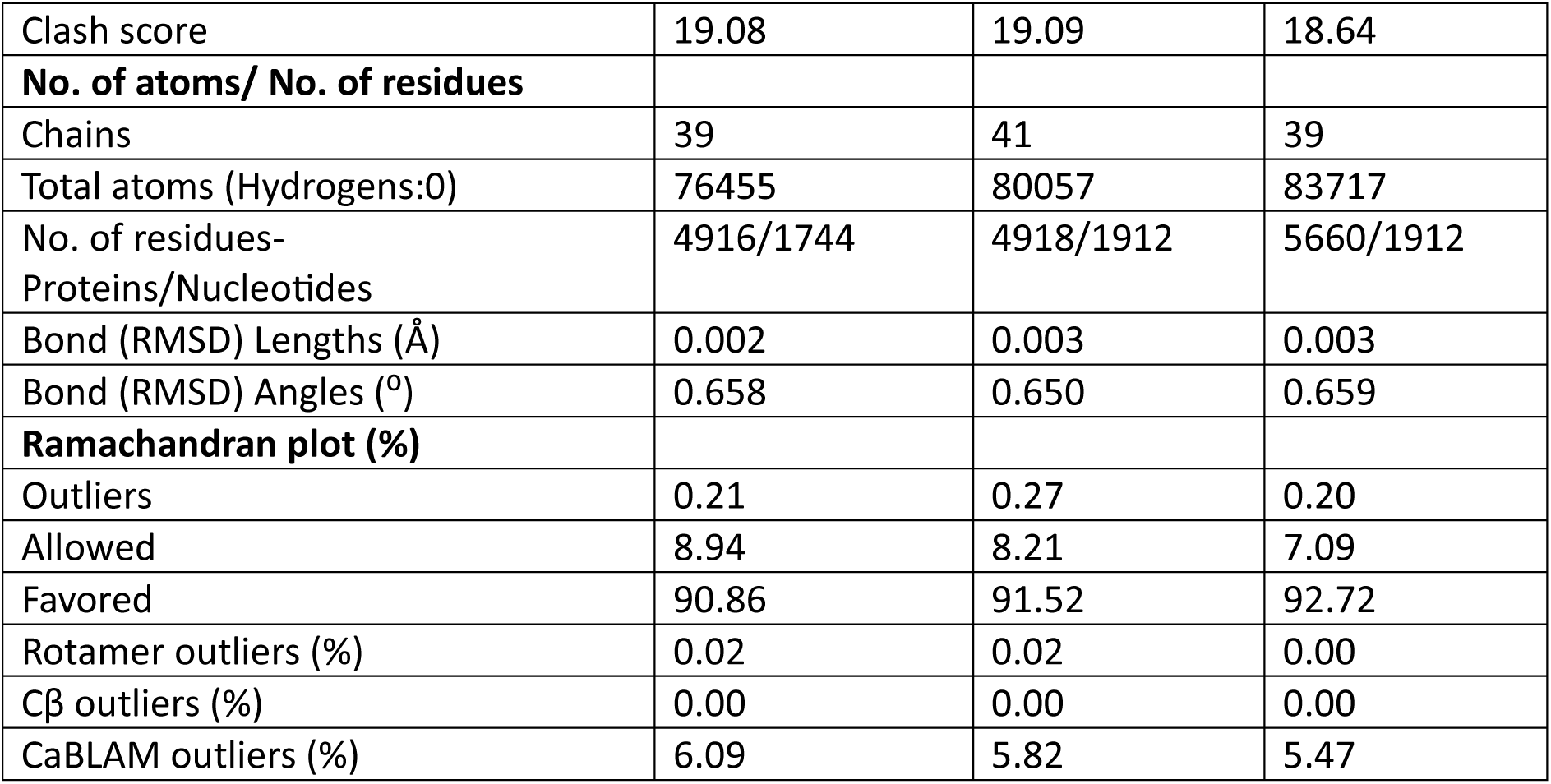
Cryo-EM data and model statistics.

### 8. Luciferase assay

WT-Luc, GNRAmut-Luc, and RAAAmut-Luc RNA was subjected to polyadenylation using *E. coli* Poly(A) Polymerase and ATP supplied in New England Biolabs kit using manufacturer’s protocol. Post polyadenylation, the RNA was extracted using RNA cleanup kit (NEB). The RNA was denatured at 95 ⁰C and refolded using buffer R at 37 ⁰C for 5 minutes. To 1 µg of RNA, 10 µl RRL was added with 0.25 µM amino acid mix minus leucine, and amino acid mix minus methionine, 1 mM ATP, 0.5 mM GTP, and the remaining volume was adjusted using buffer R to a final volume of 30 µl. Each reaction was divided into 3 sets-10 µl each and incubated at 30 ⁰C for 3 hours. The luciferase activity in each reaction was measured by adding 10 µl of Steady-Glo Luciferase reagent (Promega) and quantified using a Tecan plate reader. The graphs were plotted using Graphpad prism.

### 9. Secondary Structure Determination and Multiple Sequence Alignments

The secondary structure for I domain apex was obtained from RNAfold (*Gruber et al 2008*). Multiple sequence alignments were performed using Clustal Omega (*Sievers et al 2021*). Sequence accession number for various sequence used-EMCV (NC_001479.1), FMDV (NC_039210.1), TMEV (DQ401688.1), TRV (AB090161.1), VHEV (M80888.1), SAFV (FM207487.1), Poliovirus (NC_002058.3).

## Accession codes

Maps and atomic coordinates of the 40S ribosome (Map A), 40S ribosome-EMCV IRES-tRNA_i_ (Map B) and 40S ribosome-EMCV IRES-ternary complex (Map B1) have been deposited in the EMDB database with accession codes-EMD-64646, EMD-64644, and EMD-64645, respectively and in the PDB database with accession code-9UZM, 9UZK, and 9UZL, respectively.

## Acknowledgements

EMCV IRES (nt 280–905) into pCR2.1 was a kind gift from Dr. Bruno Sargueil, CNRS UMR8015, Universit’e Paris Descartes, France and 3C protease-pET24 was a kind gift from Prof. Raghavan Varadarajan, Indian Institute of Science (IISc), India. We thank various central facilities, namely Cryo-EM, Mass Spectrometry and Computational cluster at the Division of Biological Sciences, IISc for support. DD acknowledges the DBT-JRF Programme for fellowship. The authors acknowledge the DST-FIST support to the department. This work was supported by the Intermediate Fellowship from DBT-Welcome Trust India Alliance to TH (IA/I/17/2/503313).

