## Supplementary Figures and Tables for "Structural insights into the recruitment of viral Type 2 IRES to ribosomal preinitiation complex for protein synthesis"

***This file contains 4 supplementary figures, 2 supplementary tables and legend for supplementary movie 1.***

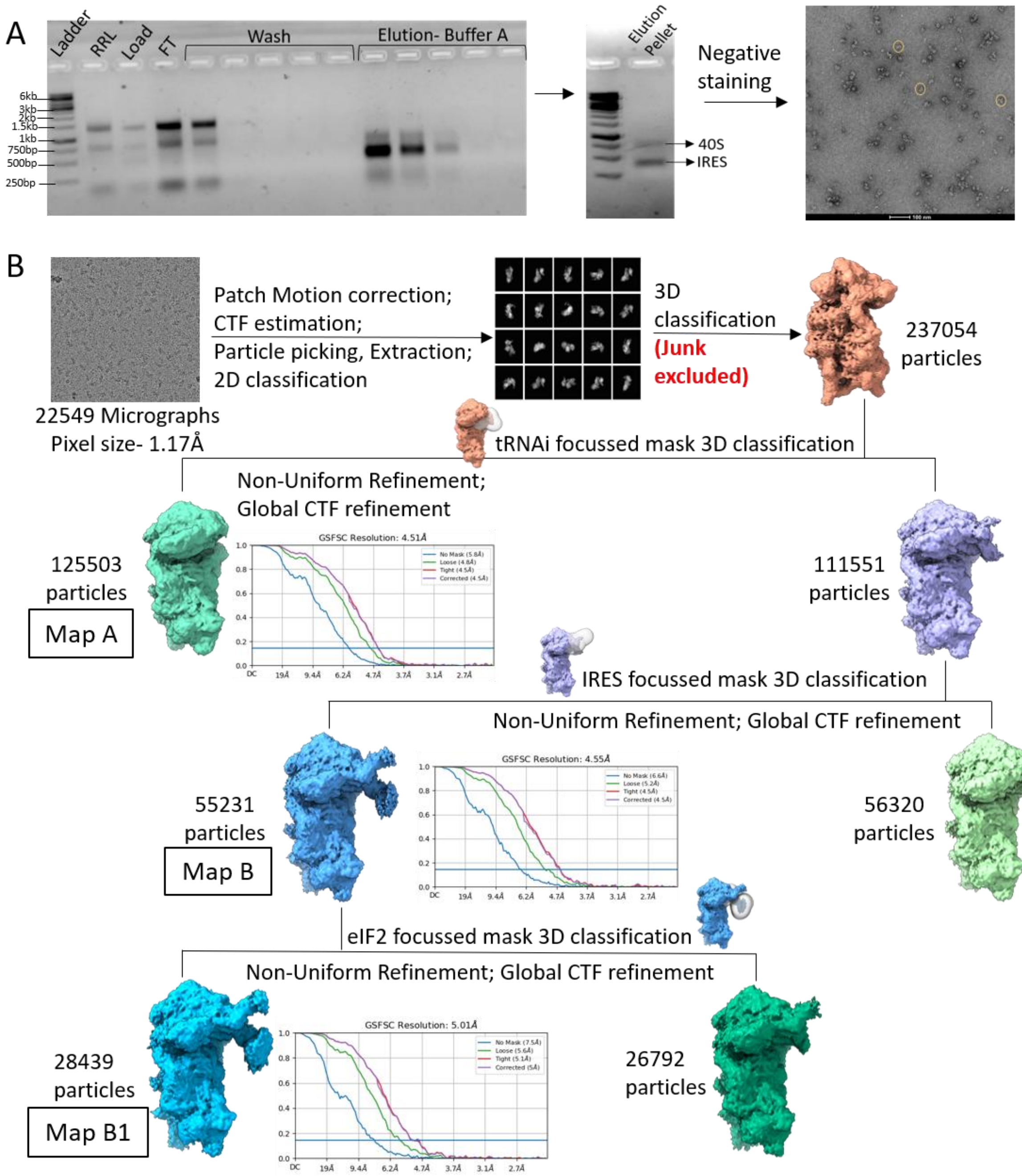


**Supplementary Figure 1.1 (A)** Isolation of EMCV IRES-48S PIC from Talon Affinity Chromatography- Talon Affinity chromatography profile and analysis of Elution fractions- pelleted by using 30% cushion on 1.5% Agarose gel, followed by Uranyl acetate staining of elution pellet and observation under transmission electron microscope at 57k X. **(B)** Processing of Cryo-EM data and features of obtained maps using CryoSPARCv4.3.0 and GSFSC resolution estimates for different maps obtained.

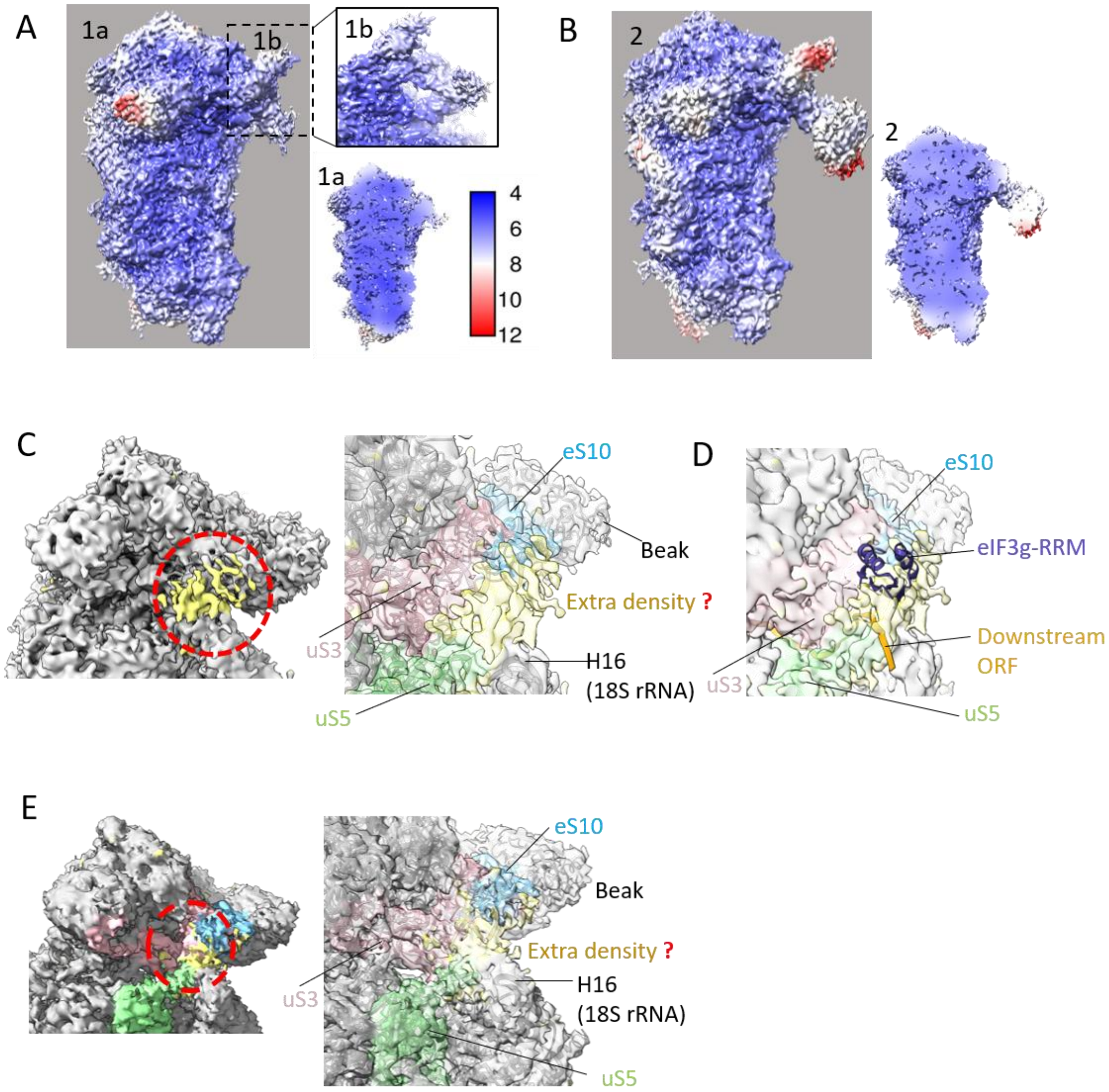


**Supplementary Figure 1.2 (A)** Local resolution estimation of Map B. (right) Surface view; (right-top) zoomed view on IRES density; (left-bottom) cross-section of Map B to show resolution at the core of the ribosome. **(B)** Local resolution estimation of Map B1. (right) Surface view; (left-bottom) cross-section of Map B1 to show resolution at the core of ribosome. **(C)** Extra density at the mRNA entry site of Map B/B1. Ribosomal proteins and RNA in contact with the unassigned extra density. **(D)** Superimposition of EMCV IRES-48S PIC on human 48S PIC (PDB Id- 8OZ0) shows the extra density coincides with the location of eIF3g-RRM and downstream mRNA. **(E)** Extra density at the mRNA entry site of Map A. Ribosomal proteins and RNA in contact with the unassigned extra density.

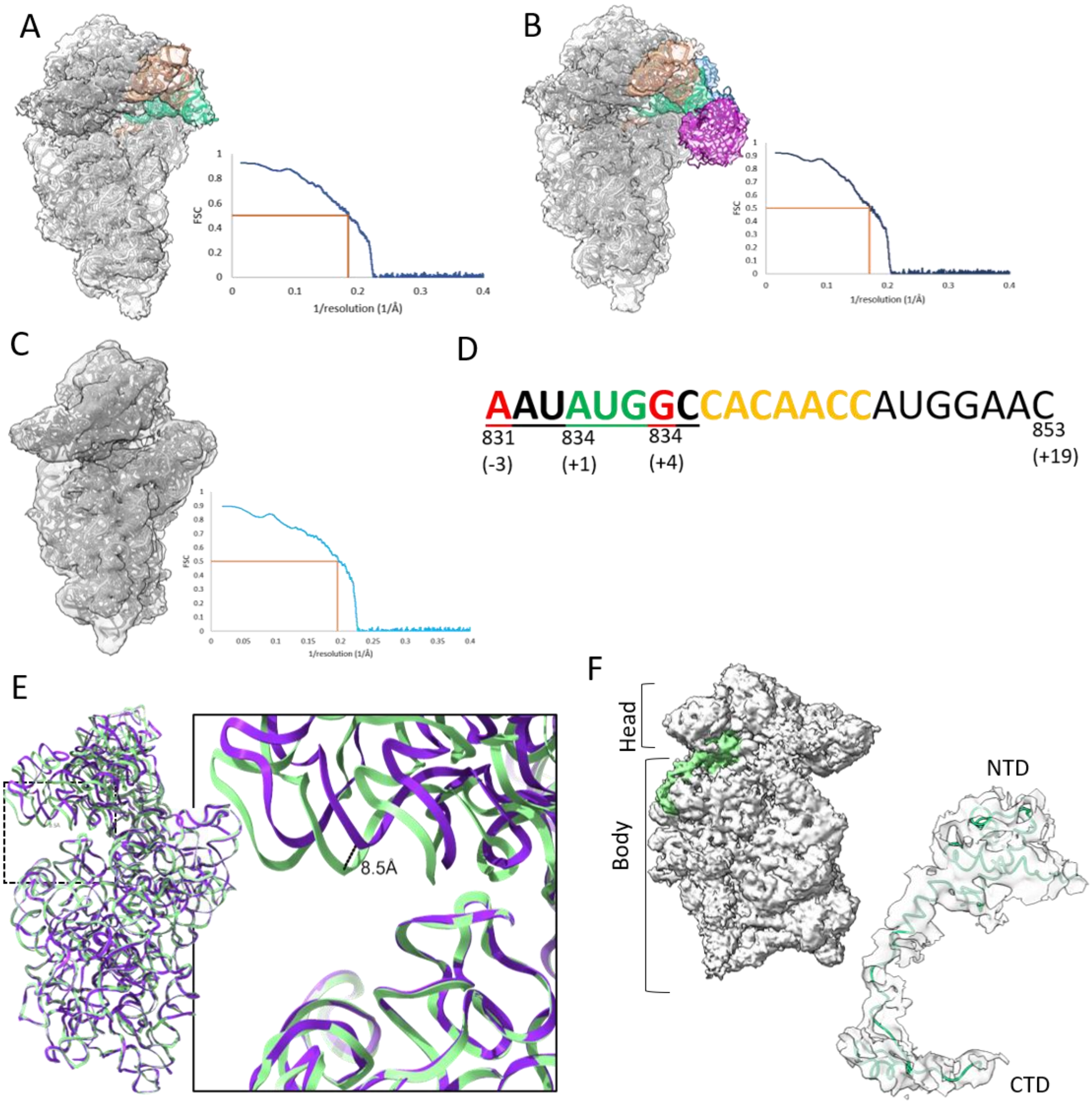


**Supplementary Figure 2 (A)** From Map B to model (EMCV IRES-48S PIC). Obtained Model showing 40S ribosome, RNA in channel, initiator tRNA, IRES domain. (right) Fourier Shell Correlation (FSC) of Map to Model fit. **(B)** From Map B1 to model (EMCV IRES-48S PIC). Obtained Model showing 40S ribosome, RNA in channel, initiator tRNA, IRES domain, eIF2αγ. (right) FSC of Map to Model fit. **(C)** From Map A to model by fitting of 40S ribosome without any factors. (right) FSC of Map to Model fit. **(D)** The sequence of EMCV IRES mRNA in the channel is provided. Residues in orange colour have no density in the obtained map B. **(E)** Comparing the state of 18S rRNA in Map A and Map B. **(F)** eS17-NTD position in 40S ribosome. (right) Fitting of eS17 in EMCV IRES-48S PIC map.

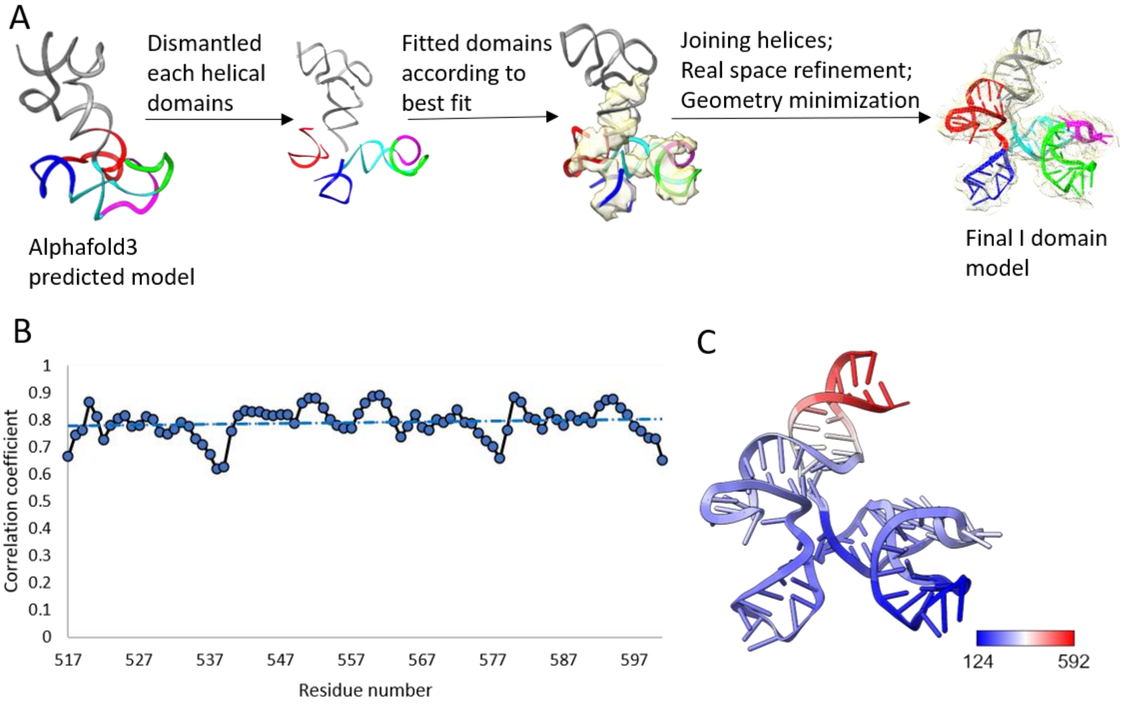


**Supplementary Figure 3.1 (A)** Process of fitting domain I apex from Alphafold3 generated model. **(B)** Correlation coefficient per residue in domain I apex for fit in the density map. **(C)** B-factor for each residue in the final model. The range is depicted in the colour key where blue is low, and red is high. **(D)** Possibility of fitting domain H in the IRES density. Domain H model was generated by Alphafold3 (ptm= 0.31). **(E)** Possibility of fitting domain J-K (PDB-8HUJ) in density. The domain J-K is way too long to fit in map and does not account for certain region. The extending loops would clash with 40S ribosome and $tRNA_i$.

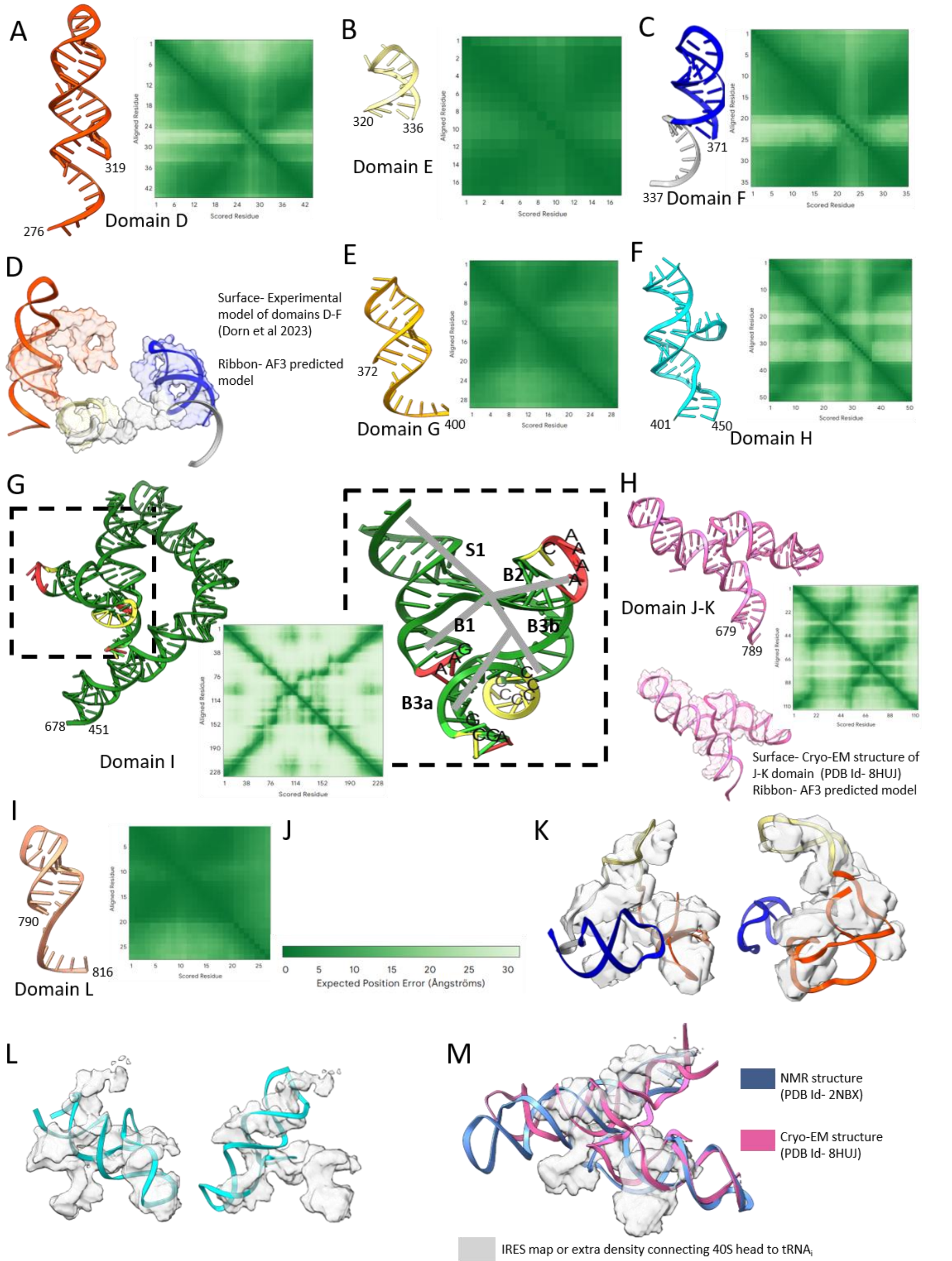


**Supplementary Figure 3.2 (A-C)** Alphafold3 predicted tertiary structures of individual domains in EMCV IRES and their corresponding predicted aligned error (PAE) plot- **(A)** domain D. **(B)** domain E. **(C)** domain F. **(D)** Fitting of predicted tertiary structure of domain D, E, F in the experimental model of domain D-F (Dorn et al 2023), showing potential correlation. **(E-G)** Alphafold3 predicted tertiary structures of domain G **(E)**, domain H **(F)**, and domain I, and zoomed view of domain I apex showing the architecture of the apical region **(G)** and its corresponding PAE plot. **(H)** Alphafold3 predicted tertiary structures of domain J-K, its PAE plot, and fitting of predicted structure in the cryo-EM model of domain J-K (PDB Id- 8HUJ). **(I)** Alphafold3 predicted tertiary structures of domain L and its PAE plot. **(J)** Color scheme describing the expected position error of the above-mentioned predictions. **(K)** Possibility of fitting domains D-F model (as determined in Dorn et al 2023) in the IRES density connecting the 40S head to tRNA in EMCV IRES-48S PIC. **(L)** Possibility of fitting domain H in the IRES density. **(M)** Possibility of fitting domain J-K (PDB- 8HUJ) in density. The domain J-K is way too long to fit in map and does not account for certain region. The extending loops would clash with 40S ribosome and $tRNA_i$.

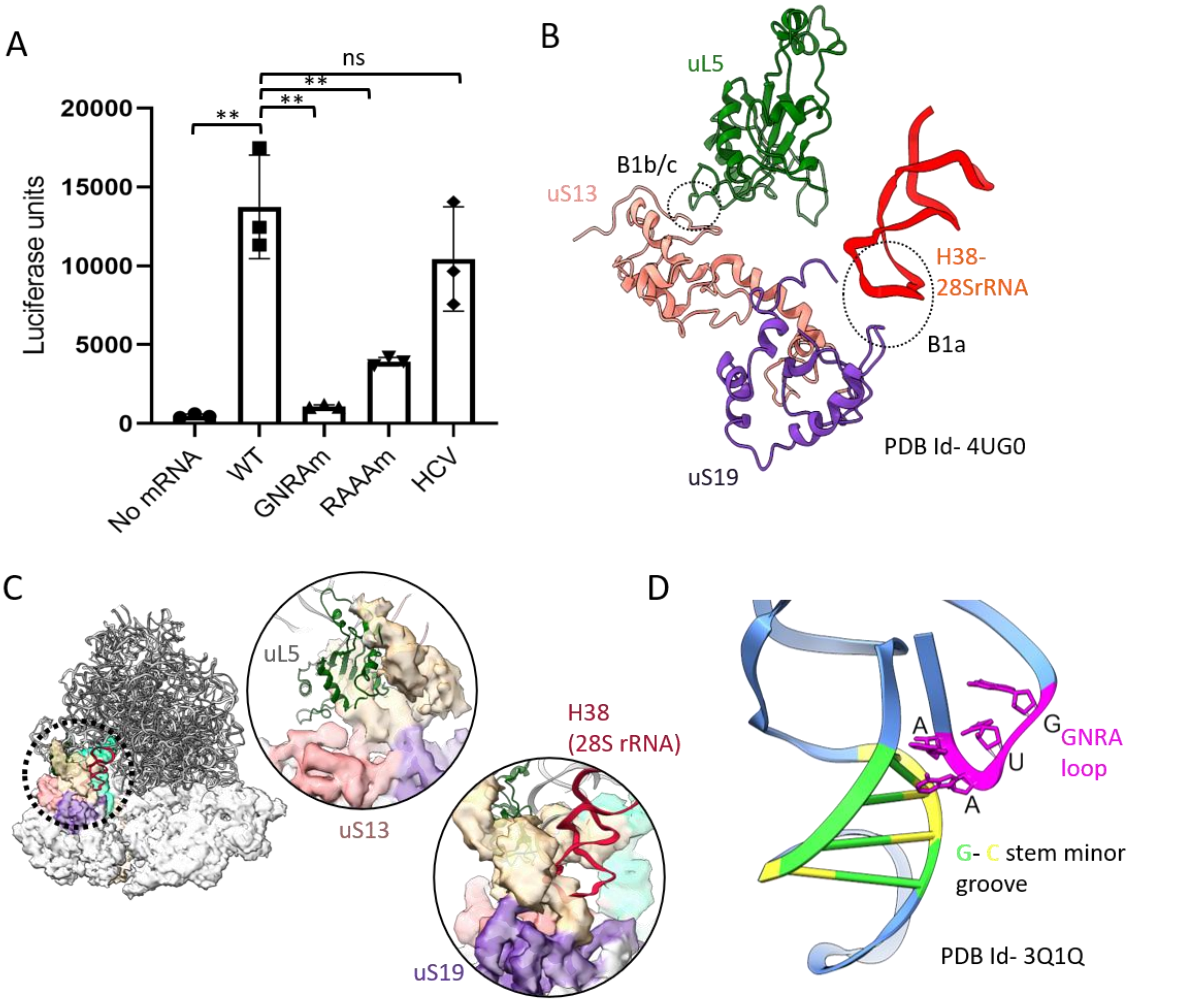


**Supplementary Figure 4 (A)** Luciferase Reporter assay showing the translation efficiency of wild type (WT) EMCV IRES with IRES mutants- GNRAm and RAAAm. The error bars represent the standard deviations of the three biological replicates. The value for each biological replicate was determined as a mean of three technical replicates. Students upaired two-tailed t test was used for statistical analysis. **(B)** Interaction of uS19 and uS13 in 40S subunit with 60S subunit- Helix 38 and uL5 to form inter-subunit bridges- B1a and B1b/c, respectively in 80S (PDB Id- 4UG0). **(C)** Estimation of clash with IRES domain I during 60S joining to form 80S-elongation competent complex (PDB Id- 4UG0). uL5 and helix 38 of 28S rRNA would clash with the IRES I domain. (left) Superimposition of 80S ribosome on EMCV IRES-48S PIC. **(D)** Reported GNRA (GUGA) interaction with the minor groove formed by C-G stem in bacterial RNaseP (PDB Id- 3Q1Q).

**Supplementary Table 1: EMCV IRES domain sequences used as inputs for Alphafold3 prediction of IRES domain tertiary structure**

| Domain | Residues | Sequence |
|---|---|---|
| D | 276-319 | CCCCCUAACGU**UACUGGCCGAAGCCGCUUGGAAUAAGGCCGGUG** |
| E | 320-336 | U**GCGUUUGUCUAUAUGU** |
| F | 337-371 | UAUUUUCCAC**CAUAUUGCCGUCUUUUGGCAAUGUG** |
| G | 372-400 | **AGGGCCCGGAAACCUGGCCCU**GUCUUCUU |
| H | 401-450 | **GACGAGCAUUCCUAGGGGUCUUUCCCCUCUCGCCAAAGGAAUGC AAGGUC** |
| I | 451-678 | **UGUUGAAUGUCGUGAAGGAAGCAGUUCCUCUGGAAGCUUCUUG AAGACAAACAACGUCUGUAGCGACCCUUUGCAGGCAGCGGAACC CCCCACCUGGCGACAGGUGCCUCUGCGGCCAAAAGCCACGUGUAU AAGAUACACCUGCAAAGGCGGCACAACCCCAGUGCCACGUUGUG AGUUGGAUAGUUGUGGAAAGAGUCAAAUGGCUCUCCUCAAGCG UAUUCAACA** |
| J-K | 679-789 | AGG**GGCUGAAGGAUGCCCAGAAGGUACCCCAUUGUAUGGGAUCU GAUCUGGGGCCUCGGUGCACAUGCUUUACAUGUGUUUAGUCGA GGUUAAAAAACGUCUAGGCC**CCCC |
| L | 790-816 | **GAACCACGGGGACGUGGUUU**UCCUUUGAAAA |

**Supplementary Table 2: List of Oligos or primers used for molecular cloning**

| Oligo name | Sequence |
|---|---|
| PTB1_Fwd | CGGGATCCATGGACGGCATTGTCCCAGA |
| PTB1_Rev | CCCAAGCTTCTAGATGGTGGACTTGGAG |
| PTB1_3C_Fwd | CGGGATCCCTGGAGGTGCTCTTCCAGGGCCCTGGCGGCTCCATGGACGGCATTGTCCCAG |
| PTB1_Stop_Rev | CCCAAGCTTCTAGATGGTGGACTTGGAGAAGG |
| EMCV_IRES_905-Fwd | CGGGATCCCCCCCTAACGTTACTGGCC |
| EMCV_IRES_905-Rev | GCTCTAGAGCAGAGCATTTTGGGCATTCCTCAAAAG |
| CAAAA_GCTGA_Rev | CAGGTGTATCTTATACACGTGGC**TCAGC**GCCGCAGAGGCACCTGTCGCCAG |
| GCGA_TACG_Rev | GCCGCAGAGGCACCTG**CGTA**CAGGTGGGGGGTTCCGCTGCCTGCAA |

**Supplementary Movie 1 Legend:**

The movie depicts the position of EMCV IRES (brown) on the 40S ribosomal subunit, in contact with $tRNA_i$ (green) in a 48S PIC state. On zooming, the IRES- domain I apical part is contacting the ribosomal proteins- uS13 (golden yellow) and uS19 (violet); elbow and acceptor arm of $tRNA_i$ which is base-paired to start codon.
